# Relationship Between Open Reading Frame 320, a Gene Causing Male Sterility in Tomatoes, and Cytoplasmic Male Sterility in Potatoes

**DOI:** 10.1101/2025.04.17.649265

**Authors:** Rika Nakajima, Rena Sanetomo, Kenta Shirasawa, Tohru Ariizumi, Kosuke Kuwabara

## Abstract

Cytoplasmic male sterility (CMS) is a phenotype wherein plants cannot develop normal male organs because of mitochondrial genes. Several cytoplasmic sterility factors are suggested to be involved in CMS phenotypes in potatoes (*Solanum tuberosum*); however, reports on the relevant mitochondrial genes remain scarce. Many potato cultivars carry mitochondrial genes that cause CMS, resulting in pollen sterility, thereby limiting their use as male parents in breeding. Therefore, identifying the causal genes is crucial for potato breeding. In this study, we focused on the T/β-type cytoplasm, the most prevalent genome type of potato worldwide, to explore mitochondrial genes involved in CMS in potatoes. We identified a novel gene, open reading frame 320 (*orf320*), in potato genotypes with T/β-type cytoplasm by comparing the mitochondrial genomes of potatoes. Functional analysis of tomatoes showed that overexpression of *orf320* with a mitochondrial transit peptide induced male sterility phenotypes, including abnormal anther development and pollen abortion. Furthermore, an investigation of *orf320* in 126 potato cultivars revealed that this gene is specific to the T/β-type cytoplasm and is absent in investigated cultivars with other cytoplasmic types. These findings provide strong evidence that *orf320* is a candidate CMS-causing gene in male sterility of T/β-type cytoplasm, offering valuable insights for future potato breeding.

## Introduction

Cytoplasmic male sterility (CMS) is a plant trait caused by the incompatibility between mitochondrial and nuclear genes, leading to abnormal male organ function. These mitochondrial genes are called CMS-causing genes (Chen and Liu 2014). CMS has been reported in over 150 plant species. Plants with CMS do not produce seeds through self-pollination; thus, CMS is a useful trait for F_1_ hybrid seed production as it eliminates the need for emasculation, a labor-intensive process. Efficient hybrid seed production using CMS plants has been implemented in various species, including rice (*Oryza sativa*), maize (*Zea mays*), sunflowers (*Helianthus annuus*), and radishes (*Raphanus sativus*) (Bohra et al. 2016). However, in potatoes (*Solanum tuberosum*), CMS is considered a major obstacle in breeding. Since potatoes are propagated vegetatively and their tubers serve as edible parts, fertile pollen production is dispensable for cultivation. However, the ability to produce fertile pollen is crucial for crossbreeding. For example, resistance genes against pests and diseases have frequently been reported in wild potato species and introduced into modern cultivars (Heřmanová et al. 2007, Vos et al. 2015). Many cultivars exhibit several different male-sterile phenotypes because of the presence of cytoplasmic sterility factors (Grun et al. 1977). Such male sterility phenotypes limit the number of potato varieties that can be used as pollen parents, thereby restricting the crossbreeding efforts of potato breeders.

In the cytoplasmic genome of potatoes, Hosaka (1986) classified chloroplast DNA (cpDNA) into five types (A, C, S, T, W), whereas Lössl et al. (1999) categorized mitochondrial DNA (mtDNA) into five types (α, β, γ, δ, ε). Hosaka and Sanetomo (2012) proposed a new nomenclature for potato cytoplasmic types (T, D, P, A, M, and W) as a rapid identification technique. These classifications are often discussed as a combination of cytoplasmic (or cpDNA) and mtDNA types. Three specific combinations—T/β (also recognized as “T” or “*tuberosum*”-type), W/α (D-type, derived from *Solanum demissum*), and W/γ—are known to associate with CMS phenotypes (Sanetomo and Gebhardt 2015). The frequency of the T/β-type cytoplasm is 72.1% in Japanese potato varieties (Hosaka and Sanetomo 2012), 59.4% in European varieties (Sanetomo and Gebhardt 2015), 73.4% in Indian varieties (Chimote et al. 2008), and 63.5% in Russian varieties (Gavrilenko et al. 2007). Thus, T/β-type cytoplasm is the most prevalent worldwide. Therefore, identifying mitochondrial genes associated with male sterility in genotypes with T/β-type cytoplasm is expected to be of great significance for potato breeding. Grun et al. (1977) suggested the existence of various cytoplasmic sterility factors ([*Sp*^*s*^], [*SM*^*s*^], [*In*^*s*^], [*TA*^*s*^], [*ASF*^*s*^], [*VSA*^*s*^], and [*Fm*^*s*^]) that cause male sterility in the presence of dominant nuclear genes (*Sp, SM, In, TA, ASF, VSA*, and *Fm*) in potatoes. These cytoplasmic factors cause various types of male sterility. However, to date, very few studies have identified the mitochondrial genes directly involved in male sterility.

In this study, we aimed to identify mitochondrial genes involved in male sterility in potato varieties with T/β-type cytoplasm and searched for genes present only in varieties with T/β-type cytoplasm, but absent in those with other cytoplasmic types, through a comparative analysis of mtDNA. We then performed gene overexpression analysis and genotyping of potato varieties and finally discussed the relationship between the identified gene and male sterility in T/β-type cytoplasm.

## Results

### Comparative analysis of mitochondrial DNA reveals open reading frames (ORFs) from genotypes with T/β-type cytoplasm

First, the mtDNA of ‘May Queen,’ the main potato variety in Japan, was constructed. The mtDNA was assembled into three contigs of 312,359, 112,802, and 50,877 bp. These mtDNA contain genes coding for NADH dehydrogenases (*nad1, nad2, nad3, nad4, nad4L, nad5, nad6, nad7*, and *nad9*), succinate dehydrogenases (*sdh3* and *sdh4*), cytochrome c reductase (*cob*), cytochrome c oxidases (*cox1, cox2*, and *cox3*), ATP synthases (*atp1, atp4, atp6, atp8*, and *atp9*), cytochrome C biogenesis group genes (*ccmB, ccmC, ccmFC*, and *ccmFN*), small subunit of ribosome (*rps1, rps3, rps4, rps10, rps12, rps13, rps14*, and *rps19*), large subunit of ribosome (*rpl2, rpl5, rpl10*, and *rpl16*), maturase (*matR*), and transport membrane protein (*mttB*). Three rRNAs (*rrn5, rrn18*, and *rrn26*) and 20 tRNAs were detected in the mtDNA (Supplementary Fig. S1).

We downloaded the mtDNAs of several cytoplasmic types listed in Supplementary Table S1 from the National Center for Biotechnology Information (NCBI) database. To identify candidate genes for T/β-type cytoplasm, we first detected ORFs encoding more than 100 amino acids in the mtDNA of T/β lines. In each genome, 160 open 263 ORFs were detected. Next, we searched for ORFs present in the mtDNA of all T/β lines and absent in the genomes of other cytoplasmic types. Finally, only three ORFs remained, i.e., *orf125, orf161*, and *orf320*, which were named based on the number of amino acids in their encoded proteins.

As most CMS-causing genes have been reported to have transmembrane domains and chimeric structures (Chen and Liu 2014), the structural characterization of the three ORFs was performed. ORF125 and ORF320, but not ORF161, contained transmembrane domains within their coding regions (Fig. 1A– C). Only ORF320 had a chimeric structure; 154 amino acids at the N-terminus of ORF320 showed 100% homology with ATP1. A transmembrane domain, which is not a chimeric sequence, was detected in the central region (Fig. 1D).

**Fig. 1.**
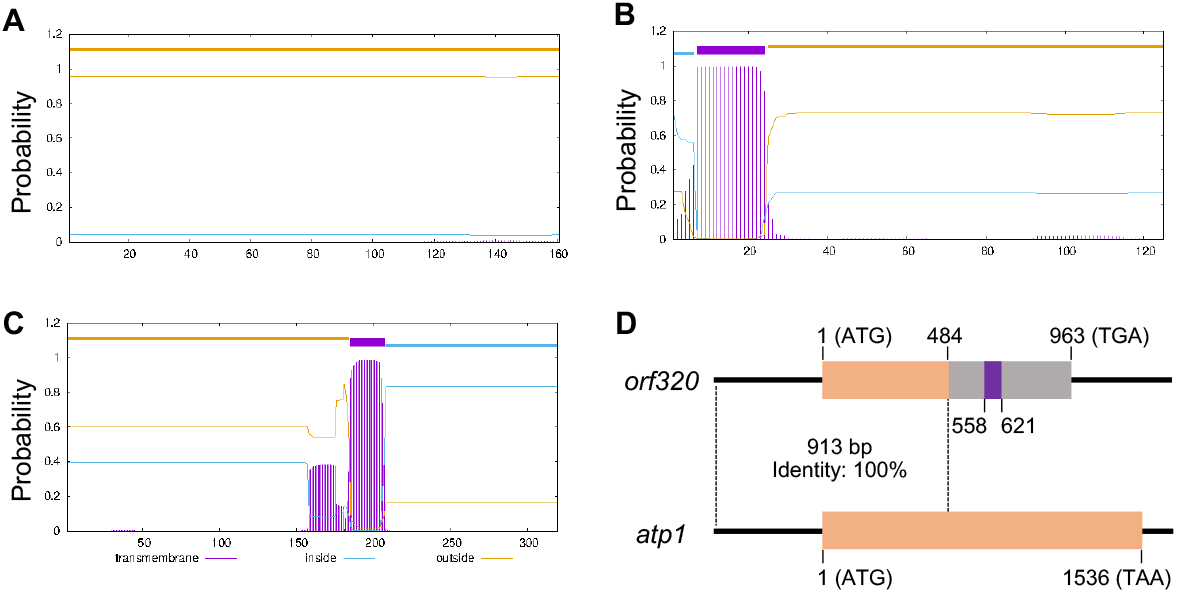
Characterization of three ORFs from potato lines with T/β-type cytoplasm. (A–C) Prediction of transmembrane domains in the three ORFs from lines with T/β-type cytoplasm. The horizontal axis indicates amino acid positions, and the vertical axis indicates the probability associated with the predicted transmembrane domain. (D) Gene structure of *orf320*. The 913 bp region surrounded by wavy lines shows 100% homology between *orf320* and *atp1*. The purple areas in *orf320* indicate the positions of its transmembrane domains.

The RNA-Seq analysis of the ‘May Queen’ cultivar revealed *orf320* expression in anthers. Reads from RNA-Seq partially mapped to the region of *orf161*, suggesting that *orf161* was weakly expressed in the anthers (Supplementary Fig. S2A–C). Moreover, reverse transcription–polymerase chain reaction (RT-PCR) analysis also revealed the expression of *orf320* in the anthers of three T/β-type potato cultivars, i.e., ‘May Queen,’ ‘Irish Cobbler,’ and ‘Kita Akari’ (Supplementary Fig. S2D). As *orf320* matched the characteristics of most CMS-causing genes reported in other plant species and was expressed in the anthers, further experiments were performed.

### Overexpression of orf320 fused with a mitochondrial-targeting peptide (MTP) causes male sterility in tomato

An overexpression vector of *orf320* was constructed and transformed into the tomato variety ‘Micro-Tom’ to investigate the association between *orf320* and male sterility. MTP was added to the N-terminus of ORF320 (Fig. 2A). In the T_0_ generation of *MTP-orf320* overexpressed lines, male sterility was observed in 12 of 33 individuals. As these individuals were male-sterile and could not produce seeds through self-pollination, we selected one *MTP-orf320* overexpressing line (*MTP-orf320-*OE83) and maintained this line by crossing with the wild-type ‘Micro-Tom’ cultivar. The sterile male phenotype was inherited stably. RT-PCR revealed the expression of *MTP*-*orf320* in the leaves, stems, roots, and anthers of *MTP-orf320-*OE83 plants (Fig. 2B). Plant growth of *MTP-orf320-*OE83 was similar to that of wild-type ‘Micro-Tom,’ suggesting that *MTP-orf320* overexpression did not affect the vegetative organs (Fig. 2C). Normal seed formation was observed when ‘Micro-Tom’ pollen was cross-pollinated with *MTP-orf320-*OE83, indicating normal female function. In contrast, abnormal anthers were observed in the *MTP-orf320-*OE83 transgenic line (Fig. 2D), suggesting that the overexpression of *MTP-orf320* severely affected male organs.

**Fig. 2.**
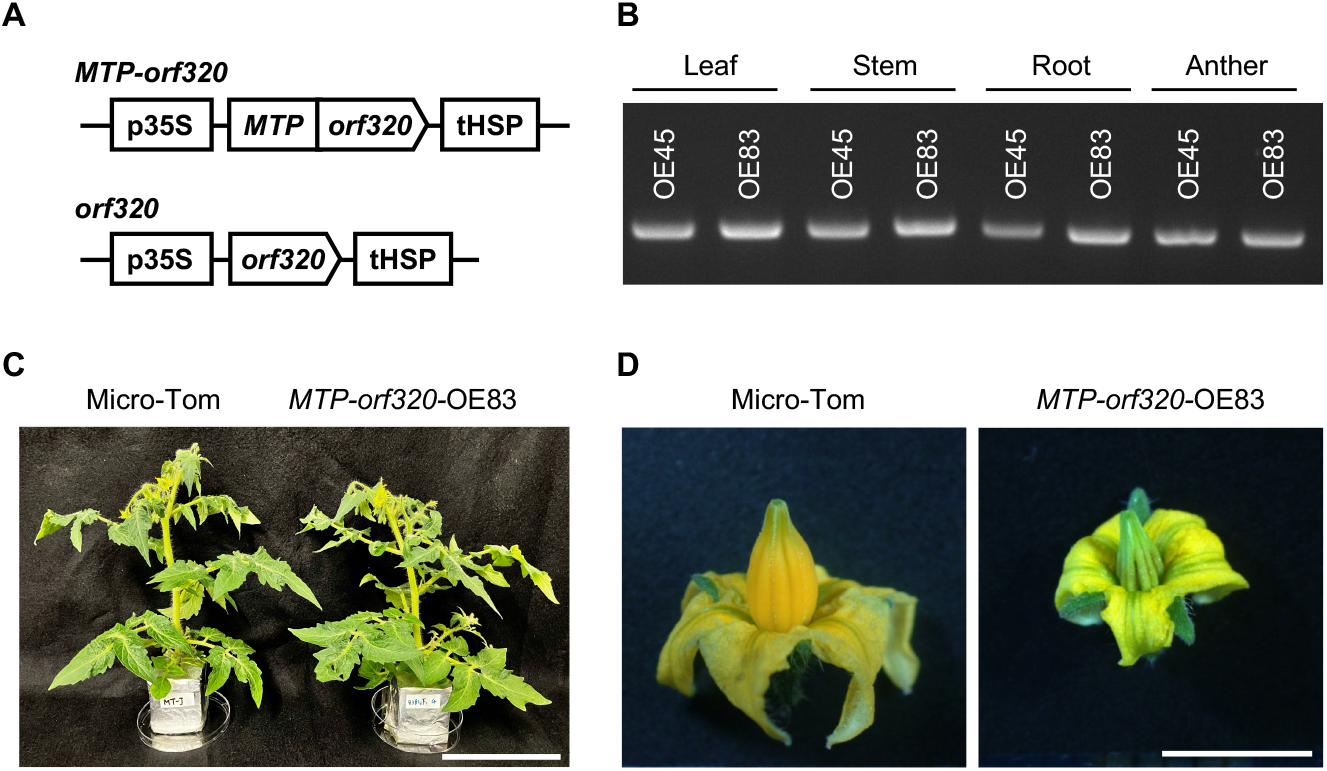
Overexpression of *orf320* induces male sterility in tomatoes. (A) Schematic illustration of the *orf320* transgene constructs used for transformation in tomato. The MTP is the N-terminal 36 amino acids of the ATPase, F1 complex, delta/epsilon subunit in *Arabidopsis thaliana* (AT5G47030). p35S, cauliflower mosaic virus 35S promoter; *MTP*, mitochondrial-targeting peptide; tHSP, heat shock protein terminator. (B) RT-PCR analysis of *MTP-orf320* in *MTP-orf320*-OE45 and *MTP-orf320*-OE83. (C) Photograph of the wild-type ‘Micro-Tom’ cultivar and the transgenic line harboring the *MTP-orf320*-OE83 transgene. Scale bar = 10 cm. (D) Photograph of flowers in the wild type ‘Micro-Tom’ cultivar and the transgenic line expressing the *MTP-orf320*-OE83 transgene. Scale bar = 5 mm.

To investigate the male-sterile phenotypes caused by *MTP-orf320* overexpression in detail, tissue sections of the anthers were observed. In the wild type, the degradation of tapetal cells gradually began from the uninucleate microspore stage and progressed during the bicellular pollen stage; the tapetal cells disappeared by the mature pollen stage (Fig. 3A). In contrast, in *MTP-orf320-*OE83, tapetal cells were similar to the wild type at the uninucleate microspore stage, but degraded prematurely and disappeared by the bicellular and mature pollen stages (Fig. 3B). These results suggest premature degradation of the tapetum layer in *MTP-orf320*-OE83.

**Fig. 3.**
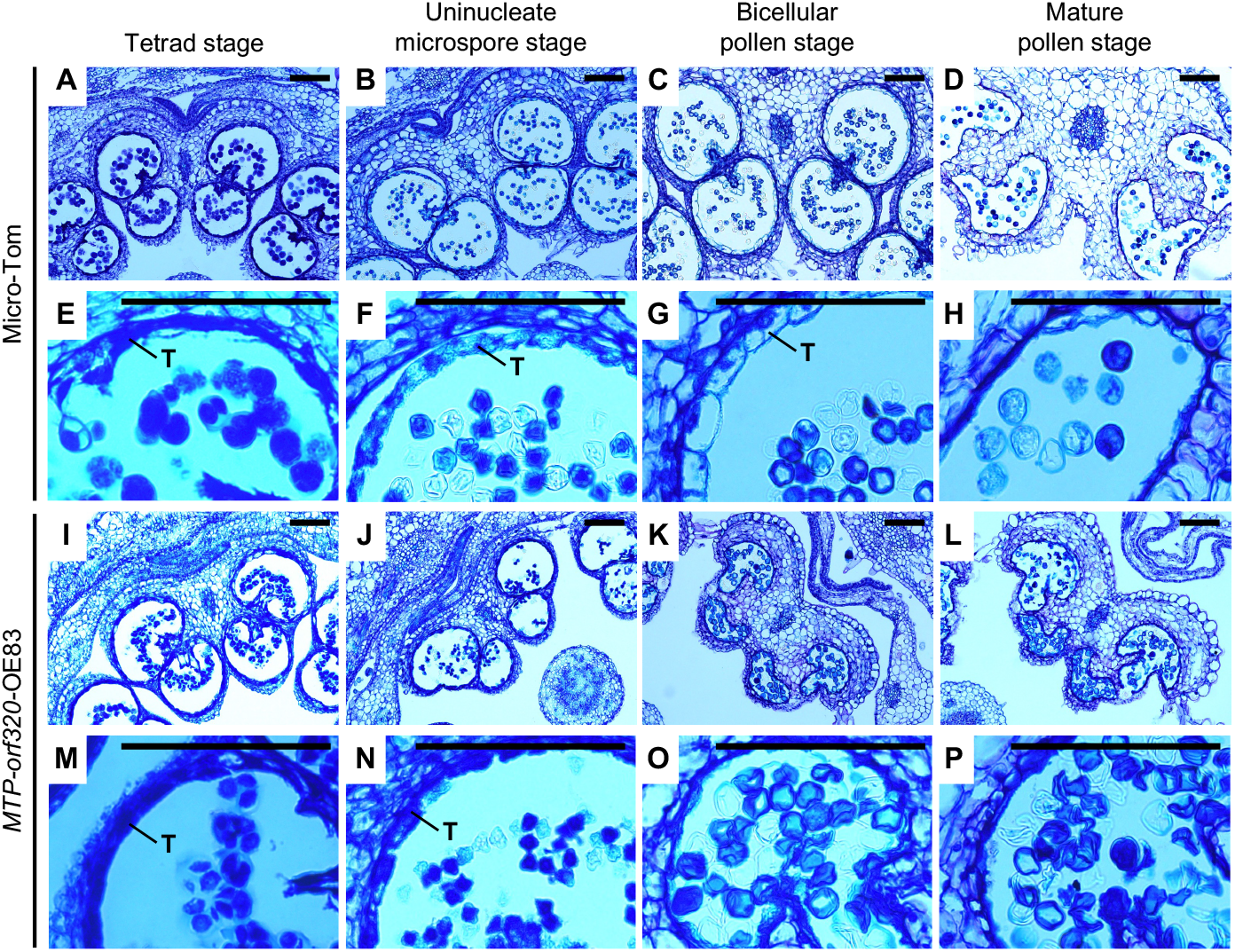
Histological examination of anthers at four different developmental stages in the wild type and *MTP-orf320* overexpressed tomatoes. (A–D) Sections of the anther tissue from the wild type ‘Micro-Tom’ cultivar and (E–H) close-up images. (I–L) Sections of the anther tissue from *MTP-orf320*-OE83 transgenic line and (M–P) close-up images. Tetrad (A, E, I, M), uninucleate microspore (B, F, J, N), bicellular pollen (C, G, K, O), and mature pollen (D, H, L, P) stages. T, tapetum layer. Scale bar = 100 µm.

As the morphology of anthers severely affects pollen development, we investigated pollen phenotypes in *MTP-orf320*-OE83 transgenic plants. In the pollen germination test, the pollen tubes were elongated in the wild type but not in *MTP-orf320*-OE83 (Fig. 4A). Alexander staining was performed to determine the viability of pollen in *MTP-orf320*-OE83 plants. Wild-type pollen was stained, whereas the pollen of *MTP-orf320*-OE83 was not (Fig. 4B). Furthermore, the number of pollen grains was significantly reduced compared with that in the wild-type (Fig. 4C). To investigate the developmental stages affected by *MTP-orf320*, we performed the DAPI (4′, 6-diamidino-2-phenylindole) staining of pollen at each stage. In the wild type, nuclei stained with DAPI were observed at the tetrad, uninucleate microspore, bicellular pollen, and mature pollen stages (Fig. 5A–D). In *MTP-orf320*-OE83, nuclei were observed at the tetrad, uninucleate microspore, and bicellular pollen stages, similar to the wild type; however, the nuclei disappeared at the mature pollen stage (Fig. 5E–H). To closely investigate the changes occurring inside the pollen from the bicellular pollen stage to the mature pollen stage, the pollen was observed using transmission electron microscopy (TEM). In the wild-type, normal development was observed from the bicellular pollen stage to the mature pollen stage, whereas in *MTP-orf320*-OE83, the vacuoles inside the pollen were enlarged at the bicellular pollen stage, indicating vacuolization within the pollen. At the mature pollen stage, fully vacuolated pollen was observed in the pollen of *MTP-orf320*-OE83 plants (Fig. 6).

**Fig. 4.**
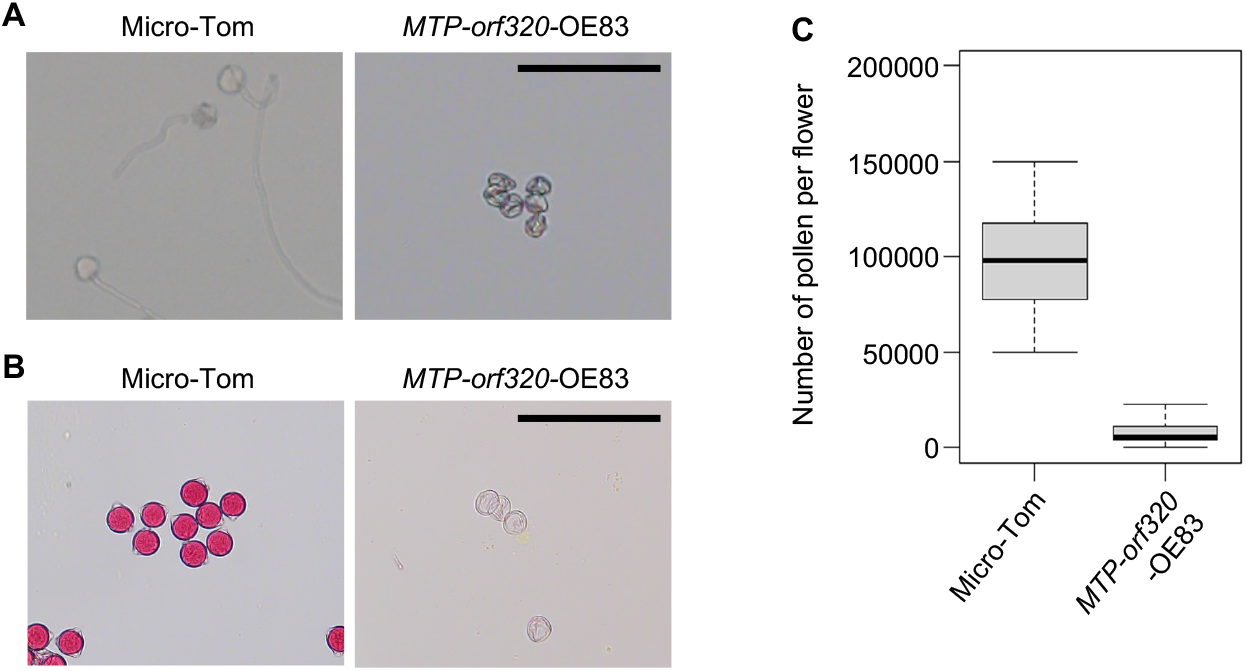
Analysis of pollen phenotypes in the wild type and *MTP-orf320* overexpressed tomatoes. (A) *In vitro* pollen germination test in ‘Micro-Tom’ and *MTP-orf320*-OE83. Scale bar = 100 µm. (B) Aborted and non-aborted pollen grains were stained by an improved Alexander staining solution. Pollen stained in red indicates fertile. Scale bar = 100 µm. (C) Number of pollen grains per flower in the wild type ‘Micro-Tom’ cultivar and *MTP-orf320*-OE83 transgenic line. The boxplots, black center lines, and whiskers represent interquartile ranges, medians, and 1.5 interquartile ranges, respectively.

**Fig. 5.**
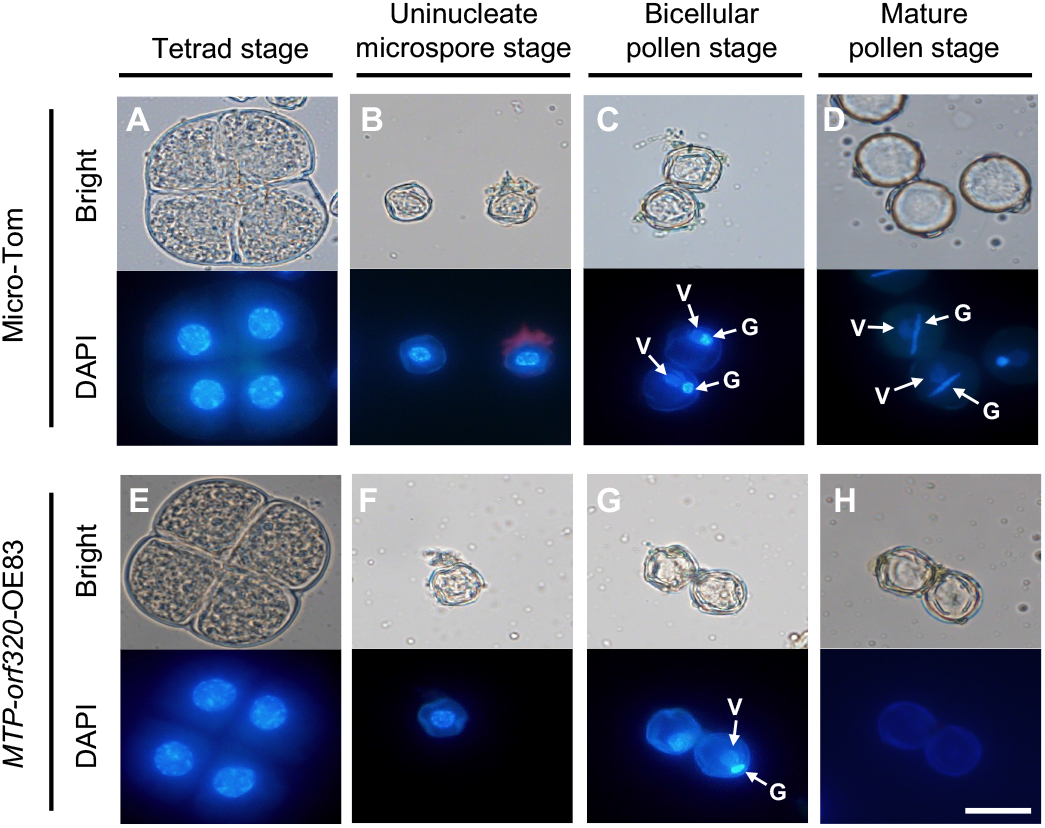
DAPI (4′, 6-diamidino-2-phenylindole) stained pollen in the wild type and *MTP-orf320* overexpressed tomatoes. (A–D) DAPI staining for the pollen of ‘Micro-Tom,’ and (E–H) *MTP-orf320*-OE83. Tetrad (A, E), uninucleate microspore (B, F), bicellular pollen (C, G), and mature pollen (D, H) stages. V, vegetative nucleus; G, generative nucleus. Scale bar = 20 µm.

**Fig. 6.**
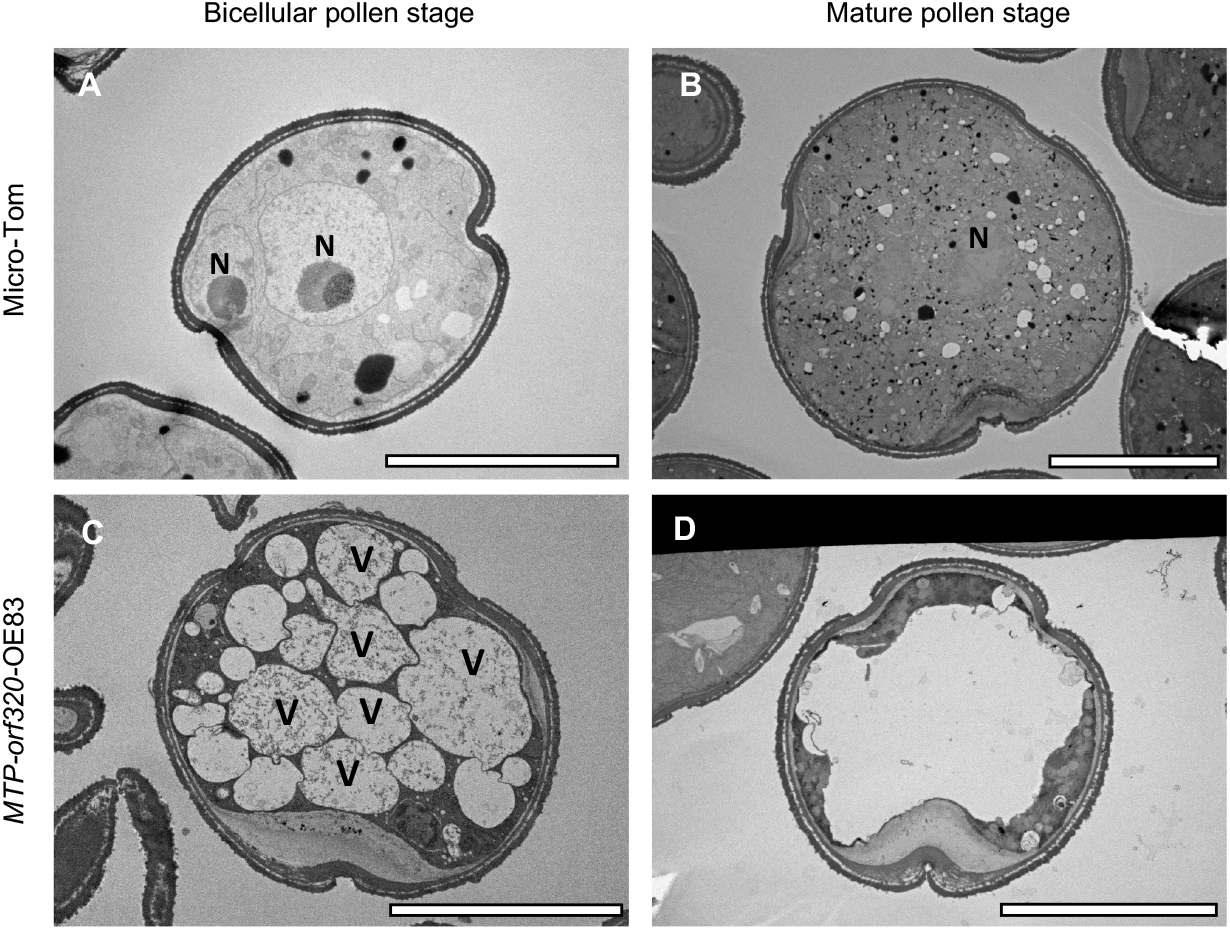
Transmission electron microscopy (TEM) analysis for pollen structure in the wild type and *MTP-orf320* overexpressed tomatoes. (A, B) TEM images of pollen structure in the wild type ‘Micro-Tom’ cultivar and (C, D) the *MTP-orf320*-OE83 transgenic line. N, nucleus; V, vacuole. Scale bar = 10 µm.

### ORF320 is transported to mitochondria without an MTP

We also transformed the *orf320* transgene construct without MTP in the ‘Micro-Tom’ cultivar. Although we expected that T_0_ plants overexpressing the *orf320* construct (without the *MTP* coding sequence) would not induce male sterility, the overexpression of *orf320* reduced fertility in 15 of 32 plants. Semi-sterile male plants had pale yellow anthers and abnormal pollen (Supplementary Fig. S3). This result suggests that *orf320* localizes to the mitochondria even without *MTP*. To investigate whether ORF320 localizes to the mitochondria without *MTP*, we generated *MTP-orf320* and *orf320* fused with *superfolder green fluorescent protein* (*sfGFP*) constructs and transiently expressed them in the protoplasts of *Nicotiana benthamiana*. The GFP fluorescence of *MTP-ORF320-sfGFP* matched that of the mitochondrial fluorescent marker MitoTracker. In the *orf320-GFP* construct, the GFP and MitoTracker fluorescence signals overlapped (Supplementary Fig. S4). These results demonstrate that ORF320 localizes to the mitochondria with and without MTP.

### Distribution of orf320 in cultivated and wild potatoes

We investigated the presence of the *orf320* gene in 126 potato cultivars with various cytoplasm types to examine whether *orf320* is specific to potato cultivars carrying T/β-type cytoplasm. *orf320* was detected in 86 out of 92 T/β-type cultivars (Table 1). The intensities of the bands obtained by PCR and electrophoresis varied among cultivars possessing *orf320* (data not shown). However, this gene was not detected in cultivars with other cytoplasm types (Table 1). This result is consistent with the results of the mtDNA comparative analysis, wherein *orf320* was detected only in the T/β-type potato lines (Supplemental Table S1), suggesting that the *orf320* gene is specific to T/β-type cytoplasm.

**Table 1.** PCR analysis for *orf320* in potato cultivars.

| Line name | cp+<br>mtDNA<br>type* | *orf320* | Line name | cp+<br>mtDNA<br>type* | *orf320* | Line name | cp+<br>mtDNA<br>type* | *orf320* |
| --- | --- | --- | --- | --- | --- | --- | --- | --- |
| Nemuromurasaki | A/ε | - | Joanna | T/β | + | Sylvia | T/β | + |
| Rankoku 3 | A/ε | - | Kanona | T/β | + | Tachibana | T/β | + |
| Harimaru | S/ε | - | Kennebec | T/β | + | Tarumae | T/β | + |
| Inca Gold | S/ε | - | Kita-akari | T/β | + | Tokachikogane | T/β | + |
| Inca-no-hitomi | S/ε | - | Kitahime | T/β | + | Tokachi Pirika | T/β | - |
| Inca-no-hoshi | S/ε | - | Kitakamui | T/β | + | Touya | T/β | + |
| Inca-no-mezame | S/ε | - | Konafubuki | T/β | + | Toyoshiro | T/β | - |
| Agasize | T/β | + | Konayuki | T/β | + | Unzen | T/β | + |
| Aino-aka | T/β | + | Rirachip | T/β | + | Waseshiro | T/β | + |
| Aiyutaka | T/β | + | Matilda | T/β | + | White Flyer | T/β | - |
| Akane Kaze | T/β | + | May Queen | T/β | + | Yankee Chipper | T/β | + |
| Alowa | T/β | + | May Queen | T/β | + | Yellow Shark | T/β | + |
| Andover | T/β | + | Myojo | T/β | + | Yukijiro | T/β | + |
| Atlantic | T/β | + | Neodelicious | T/β | + | Yukirasha | T/β | + |
| Beni-akari | T/β | + | Niseko | T/β | + | Yukitsubura | T/β | + |
| Benimaru | T/β | - | Nishiyutaka | T/β | + | Andigena 553-4 | W/α | - |
| Bifuka Kurenai | T/β | + | Norin 1 | T/β | + | Astarte | W/α | - |
| Bihoro | T/β | + | Norin 2 | T/β | + | Benihisashi | W/α | - |
| Chelsea | T/β | + | Norking Russet | T/β | + | Chijiwa | W/α | - |
| Chipeta | T/β | + | North Chip | T/β | + | Eva | W/α | - |
| Chitose | T/β | + | Oojiro | T/β | + | Ezo-akari | W/α | - |
| Corolle | T/β | + | Okhotsk-chip | T/β | + | Hanashibetsu | W/α | - |
| Cynthia | T/β | - | Pike | T/β | + | Kitamurasaki | W/α | - |
| Danshaku | T/β | + | Pike | T/β | + | Koganemaru | W/α | - |
| Dejima | T/β | + | Poroshiri | T/β | + | Konayutaka | W/α | - |
| Desiree | T/β | + | Prevalent | T/β | + | Meihou | W/α | - |
| Dorothy | T/β | + | Ranran-chip | T/β | + | Monticello | W/α | - |
| Dr. Johansen | T/β | - | Red Moon | T/β | + | Musamaru | W/α | - |
| Early Starch | T/β | + | Russet Burbank | T/β | + | Natsufubuki | W/α | - |
| Eniwa | T/β | + | Sakurafubuki | T/β | + | Northern Ruby | W/α | - |
| Fugenmaru | T/β | + | Sanjoomaru | T/β | + | Pearl Starch | W/α | - |
| Ground Pechka | T/β | + | Saya-akane | T/β | + | Pirka | W/α | - |
| Haru-akari | T/β | + | Sayaka | T/β | + | Rishiri | W/α | - |
| Haruka | T/β | + | Shepody | T/β | + | Sassy | W/α | - |
| Hatsufubuki | T/β | + | Sherry | T/β | + | Setoyutaka | W/α | - |
| Hikaru | T/β | + | Shimabara | T/β | + | Shadow Queen | W/α | - |
| Hokkai-aka | T/β | + | Shizuki | T/β | + | Shiretoko | W/α | - |
| Hokkaikogane | T/β | + | Salem | T/β | + | Tawaramurasaki | W/α | - |
| Inca Purple | T/β | + | Snow March | T/β | + | Toyo-akari | W/α | - |
| Inca Red | T/β | + | Snowden | T/β | + | Tunika | W/α | - |
| Innovator | T/β | + | Star Queen | T/β | + | Youraku | W/α | - |
| Jaga Kids Purple | T/β | + | Star Ruby | T/β | + | Konahime | W/γ | - |

+ indicates that the gene was confirmed to be present by PCR.
- indicates that the gene was confirmed to be absent by PCR.
\* indicates the cytoplasm types reported in Hosaka and Sanetomo (2012).

In addition, 559 accessions of wild species, and 290 accessions of cultivated species, were investigated for the presence and geographical distribution of *orf320*. Wild potato species harboring *orf320* were frequently detected in the southern Andes, particularly in Bolivia and Argentina (Supplementary Fig. S5). *orf320* was not detected in Andean-cultivated species (*S. tuberosum* ssp. *andigena*) with cp DNA types A, C, and S; however, it was consistently detected in common potatoes (*S. tuberosum* ssp. *tuberosum*) with T-type cytoplasm. However, as *orf320* has been detected in many wild species (Supplementary Table S2), its origin and transmission to *S. tuberosum* ssp. *tuberosum* remains unclear.

## Discussion

### orf320 has a chimeric structure and a transmembrane domain

We compared the mtDNAs of different cytoplasmic types and detected three ORFs (*orf125, orf161*, and *orf320*) in genotypes with T/β-type cytoplasm. Most CMS-causing genes identified in plant species have transmembrane domains and chimeric structures (Chen and Liu 2014). Of ORFs from T/β-type cytoplasm, only *orf320* shows characteristics consistent with many CMS-causing genes. *orf320* formed a chimeric structure with *ATP1* at N-terminal 154 amino acid (Fig. 1D). It has been previously reported that *orf79*, a CMS-causing gene in Boro-Taichung 65 type cytoplasmic male sterile rice (BT-CMS), has a chimeric structure with *cox1* and an unknown sequence (Kazama et al. 2008). In rice with wild-type abortive-type CMS (WA-CMS), the CMS-causing gene *WA352* exhibits a chimeric structure composed of three genes, *orf284, orf224*, and *orf288*, with unknown functions. A portion of *orf288* in WA352 interacts with the nuclear-encoded mitochondrial protein COX11 and triggers premature tapetal programmed cell death (PCD) in the anther tapetum (Luo et al. 2013). ORF320 also contained a transmembrane domain in the central region, which was not represented by a chimeric sequence (Fig. 1D). This property is common in most CMS-causing genes. For example, URF13 in maize CMS-T (Texas), which has a transmembrane domain, has been reported to form pores in the inner membrane (Rhoads et al. 1995). Based on a report on URF13, it has been argued that CMS-causing genes with transmembrane domains disrupt the proton gradient in the inner mitochondrial membrane and affect ATP synthesis (Chen and Liu 2014). The transmembrane domain in the coding region of ORF320 may be involved in disrupting the proton gradient in the inner membrane.

### orf320 derived from potato causes male sterility in tomato

Most CMS-causing genes induce male sterility in plant species. *WA352* in rice WA-CMS causes male sterility when overexpressed in *Arabidopsis thaliana* with a mitochondrial transit peptide (Luo et al. 2013). *orf147*, discovered in pigeon peas (*Cajanus cajanifolius*), causes male-sterile phenotypes when overexpressed in *Arabidopsis* via a mitochondrial transition signal (Bhatnagar-Mathur et al. 2018). The CMS-causing gene *orf125* in *Raphanus sativus* was introduced into the mitochondrial genome of *Brassica napus* by cell fusion, and the cybrid exhibited a CMS phenotype (Arimura et al. 2018, Sakai and Imamura 1992). In this study, *orf320* from T/β-type potatoes was overexpressed in tomatoes, resulting in male sterility (Fig. 2D). As most CMS-causing genes induce male sterility in different plant species, *orf320* is a promising candidate for a CMS-causing gene in potatoes.

### Overexpression of orf320 causes the premature degradation of tapetum and pollen abortion

The overexpression of *orf320* with MTP under the control of 35S promoter causes severe male-sterile phenotypes in tomatoes. The histological analysis of the anthers revealed premature tapetum degradation in *MTP-orf320* overexpressing lines (Fig. 3). Anthers play an important role in microspore development by secreting proteins, lipids, and enzymes. Subsequently, the tapetum undergoes timely PCD regulated by reactive oxygen species (ROS) and cyt c release. Disorders in these components lead to male sterility (Biswas and Chaudhuri 2024). It has been reported that mitochondrial CMS-causing genes are related to the disruption of ROS and Cyt c. Overexpression of *orf507*, a CMS-causing gene in pepper (*Capsicum annuum*), under a tapetum-specific promoter, leads to an increase in the activity of cytochrome c oxidase, ultimately inducing premature PCD of the tapetum layer (Ji et al. 2015). WA352 in rice WA-CMS inhibits the function of COX11 in peroxide metabolism, leading to massive ROS and Cyt c release, eventually causing premature tapetal PCD and male sterility (Luo et al. 2013). Based on these reports, *orf320* may induce massive ROS and Cyt c release in the anther tapetum, leading to premature PCD. We also performed a phenotypic analysis of the pollen and found that the overexpression of *MTP-orf320* caused abnormal pollen phenotypes, such as non-germination and non-staining (Fig. 4). Because pollen development depends completely on tapetum degradation (Biswas and Chaudhuri 2024), premature tapetum degradation in *MTP-orf320* overexpressed lines may cause pollen abortion.

Notably, reduced fertility was observed when *orf320* was expressed without MTP (Supplementary Fig. S3). A similar phenomenon was reported for the overexpression of *orf288*, a CMS-causing gene identified in *Brassica juncea*. When *orf288* was expressed in *Arabidopsis* with or without a mitochondrial transit peptide, both constructs showed sterile male phenotypes. Furthermore, it has been confirmed that ORF288 localizes to mitochondria without the assistance of a transit peptide (Jing et al. 2012). Similarly, ORF320 was found to localize to the mitochondria without MTP (Supplementary Fig. S4). Considering the subcellular localization results, it is suggested that ORF320 induces male sterility by localizing and functioning in the mitochondria.

### orf320 is likely associated with male sterility in T/β-type cytoplasm

At least seven cytoplasmic sterility factors cause different male-sterile phenotypes in potatoes (Grun et al. 1977). In addition, frequently reported male sterile phenotypes in T/β-type potato cultivars are abnormal anther morphology, reduced pollen number, and low pollen staining rate (Santayana et al. 2022). Notably, these phenotypes closely resembled the male sterility observed in *orf320*-overexpressed tomato (*MTP-orf320*-OE83), showing abnormal anther development, reduced pollen number, and unstained pollen (Figs. 2–6). Furthermore, *orf320* was detected in most potato cultivars with T/β-type cytoplasm and not in cultivars with other cytoplasm (Table 1). These results suggest that *orf320* is associated with male sterility in the T/β-type cytoplasm and may be one of the cytoplasmic sterility factors previously reported by Grun et al. (1977). Additionally, the band intensities of the PCR-amplified fragments of the *orf320* gene varied among the cultivars wherein it was detected, suggesting that the copy number of *orf320* may differ among cultivars. Even in the six cultivars judged to have no bands, it is possible that they retained *orf320* at very low copy numbers. However, the relationship between the copy number of this gene and CMS expression remains unclear. Further studies are required to confirm this association.

However, a gene predicted to encode another cytoplasmic factor has been recently reported. In the male sterile somatic hybrids between common potato (*S. tuberosum*) and wild species *S. commersonii, orf125* was identified as a CMS-causing gene and participated in the onset of “*tuberosum*”-type, i.e., T/β-type CMS (Tamburino et al. 2024). Notably, *orf125* was detected in our mtDNA comparative analysis (Fig. 1; Supplementary Table S1), supporting the validity of the analytical method used in this study.

To demonstrate that *orf320* is the gene responsible for male sterility in T/β-type cytoplasm, two strategies are required. In other plant species, the transcripts of CMS-causing genes are regulated by the restorer-of-fertility genes (Chen and Liu 2014). A few cultivated potato varieties, such as ‘Kita-akari,’ exhibit pollen fertility despite carrying T/β-type cytoplasm. Iwanaga et al. (1991) also reported the presence of a dominant restoration gene in *S. tuberosum*. Therefore, it is necessary to investigate whether *orf320* is transcriptionally regulated in potato cultivars that exhibit pollen fertility. In recent years, the knockout of mitochondrial genes using transcription activator-like effector nuclease-mediated mitochondrial genome editing (mitoTALEN) has been reported in various plant species, such as *Arabidopsis*, rice, rapeseed (*Brassica napus*), tomato, and potato (Ayabe et al. 2023, Kazama et al. 2019, Kuwabara et al. 2022, Nicolia et al. 2024). This approach enables the targeted disruption of mitochondrial genes to assess their functional roles. To further validate the involvement of the *orf320* gene in CMS, it is essential to examine whether knocking out this gene in male-sterile cultivars, such as ‘May Queen,’ restores pollen fertility. The use of these two strategies is expected to provide critical insights into the molecular mechanisms underlying CMS in potatoes and promote potato breeding in the future.

## Materials and Methods

### Plant materials

A miniature dwarf tomato cultivar, ‘Micro-Tom’ (Scott et al. 1989), was cultivated at 23°C under a 16/8 h light/dark cycle. The potato cultivars, ‘May Queen,’ ‘Irish Cobbler,’ and ‘Kita Akari,’ which possess T/β-type cytoplasm, were grown in a greenhouse. DNA from 126 potato cultivars, 559 accessions of wild species, and 290 accessions of cultivated species used for screening of *orf320* was preserved at the Potato Germplasm Enhancement Laboratory of Obihiro University of Agriculture and Veterinary Medicine. The Chloroplast type of these materials was determined by Sukhotu and Hosaka (2006).

### Genome sequence analyses

Total genomic DNA was extracted from the young leaves of ‘May Queen’ with a Maxwell 16 Instrument and Maxwell 16 Tissue DNA Purification Kits (Promega). A HiFi SMRTbell library was constructed using the SMRTbell Express Template Prep Kit (version 2.0; PacBio). The resulting library was separated using BluePippin (Sage Science) to remove short DNA fragments (<15 kb) and sequenced using 8 M SMRT cells on a PacBio Sequel II system (PacBio).

### Genome assembly and annotation

Using DNA purified from the leaves of the potato cultivar May Queen, 150 bp paired-end sequence data were obtained using the short-read Illumina NovaSeq 6000. These sequences were trimmed using the “Trim Reads” tool in Genomic Workbench ver. 23.05 with the parameters set to *p* = 0.05 and a minimum length of 50. The trimmed sequences were mapped to the previously reported mtDNA of the potato cultivar Desiree (NCBI accession numbers MN104801, MN104802, and MN104803), and a consensus sequence was obtained. Next, the ‘May Queen’ sequence data generated with the long-read sequencer PacBio Sequel II were mapped to this consensus sequence to construct longer contigs. Thus, regions with low coverage appeared as “N” in the sequence. These regions were manually inspected using mapping data and corrected, leading to the reconstruction of the ‘May Queen’ mtDNA. The assembled genome was annotated using the online tool PMGA with Dataset 2 (Li et al. 2024) and visualized using OGDRAW (Greiner et al. 2019).

### ORF analyses

The complete mtDNA sequences are listed in Supplementary Table S1 and were downloaded from the NCBI database. Initially, ORFs encoding more than 300 nucleotides (≥100 amino acids) on the mtDNA of T/β-type potato lines were detected using getorf (https://emboss.sourceforge.net/apps/cvs/emboss/apps/getorf.html; Version: EMBOSS:6.6.0.0). ORFs were compared against the mtDNA of other than T/β-type potato lines using BLASTN (Ye et al. 2006) and parameter settings “-evalue 1e-5 -perc_identity 99 -qcov_hsp_perc 99.” BLASTN obtained common ORFs between T/β-type potato lines with 100% of sequence homology. Transmembrane domains in the coding sequences were predicted using TMHMM-2.0 (Krogh et al. 2001).

### Vector construction and expression of orf320 in tomato

The *p35S-MIR-tHSP* vector forms the backbone of the reconstructed vector, which contains the cauliflower mosaic virus 35S promoter, miraculin gene (*MIR*), and heat shock protein (*HSP*) gene terminator (Ono et al. 2021). The *MIR* sequences of the vectors were replaced with *MTP-orf320* or *orf320. MTP, orf320*, and the vector backbone were amplified using Prime STAR GXL Premix (Takara Bio) and Prime STAR GXL Premix Fast Dye Plus (Takara Bio). Gel extraction of the PCR products was performed using the Fast Gene Gel/PCR Extraction Kit (Nippon Genetics) following the manufacturer’s protocol. Subsequent In-Fusion cloning was performed using the In-Fusion HD Cloning Kit (Takara Bio) to create a transformation vector. The resulting transformation vector was introduced into *Agrobacterium* strain GV3101. Transformation of the tomato cultivar ‘Micro-Tom’ was performed according to the method described by Sun et al. (2006) with kanamycin selection. The T_0_ plants were selected by PCR using primers for *neomycin phosphotransferase II* (*NPTII*; Supplemental Table S3).

### RNA expression analyses

Total RNA was extracted from the anthers of potato cultivars and transgenic tomatoes using an RNeasy Plant Mini Kit (Qiagen, Hilden, Germany). The extracted RNA was treated with RNase-free DNase (QIAGEN) and used for sequence library preparation using the TruSeq Stranded mRNA Library Prep Kit (Illumina), in which random hexamers were used as primers for reverse transcription. The resultant library was sequenced on a DNBSEQ G400 instrument (MGI Tech) to generate 100-bp paired-end reads.

Adaptors and low-quality reads were trimmed using Trim_galore (https://github.com/FelixKrueger/TrimGalore) with option -q 30. Hisat2 (Kim et al. 2015) was used for mapping reads onto the assembled genome of ‘May Queen,’ and Integrative Genomics Viewer (Robinson et al. 2011) was used for visualization.

RT-PCR was performed to validate RNA-Seq results. In total, 800 ng of total RNA isolated from the tissues of transgenic tomatoes was converted into cDNA using the ReverTra Ace qPCR RT Kit (TOYOBO) with random primers (Takara Bio). The PCR mix (10 μL) contained 0.5 μL cDNA, 0.5 μM primers (Supplemental Table S3), and 5 µL GoTaq Green Master Mix (Promega). The thermal cycling conditions were as follows: initial denaturation at 95°C for 2 min; 35 cycles of denaturation at 95°C for 30 s, annealing at 58°C for 30 s, and extension at 72°C for 60 s; and a final extension at 72°C for 5 min. PCR products were separated by electrophoresis on a 1% agarose gel in Tris-acetate-EDTA buffer. The gels were stained with Midori Green Advance (NIPPON Genetics) to detect the DNA bands under ultraviolet illumination.

### Histological analysis of anthers

The histological analysis of the anthers was performed as described by Hao et al. (2017). Paraffin-embedded anthers were sectioned into 10 µm thickness using a MICROM HM325 microtome (PHC). The sections were placed on microscope slides and dried overnight at 42°C. The sections were stained by immersion in 0.05% toluidine blue for 5 min, gently rinsed with water, and dried. The slides were immersed in xylene for a minute to remove the paraffin. Before drying the xylene, a drop of Entellan New (Merck Millipore) was applied to the microscope slides, which were then covered with a coverglass and left to stand overnight. Sections were observed and photographed using an OLYMPUS BX53 optical microscope (OLYMPUS).

### Phenotypic analysis of pollen

The *in vitro* pollen germination tests were performed following a previously described method (Kuwabara et al. 2022). To count the total number of pollen grains, flowers at the anthesis stage were immersed in water and vortexed to release the pollen. Pollen suspensions were input into a cell counter (Biomedical Science), and the total number of pollen grains was calculated according to the manufacturer’s instructions. The aborted and non-aborted pollen grains released from the anthers during anthesis were stained using an improved Alexander staining solution (Peterson et al. 2010). For DAPI staining, pollen or microspores from the anthers at the pollen developmental stage were fixed in EAA (EtOH: acetic acid = 3:1). Subsequently, pollen or microspores were stained using DAPI staining solution, containing 0.1 M phosphate buffer (pH 7.0), 0.5 M EDTA, Triton X-100, and 3.0 µg/mL DAPI, for more than 5 min and observed using ultraviolet light under the OLYMPUS BX53 optical microscope. To analyze pollen structure using TEM, pollen samples collected from the anthers after flowering were sampled into a fixative solution, containing 2% paraformaldehyde, 2% glutaraldehyde, and 0.05 M cacodylate buffer (pH 7.4), and fixed by chilling at 4°C overnight. The pollen was then washed by replacing it with 0.05 M cacodylate buffer, allowed to stand for 30 min, and repeated three times. The pollen was replaced with 2% osmium tetroxide (OsO_4_) in 0.05 M cacodylate buffer and kept overnight at 4°C. The pollen was then dehydrated through a graded ethanol series of 50 and 70% ethanol at 4°C for 60 min each, followed by dehydration in 90% ethanol for 60 min at 25°C. Subsequently, the pollen was dehydrated three times for 60 min each with 100% ethanol at 25°C and then left in 100% ethanol for 2 days for complete dehydration. The pollen was then replaced with propylene oxide and left for 30 min; this process was repeated twice. Next, the pollen was left to stand in a propylene oxide: resin solution (50:50 v/v, Quetol-651; Nisshin) for 6 h. The pollen was then incubated for 16 h in 100% resin. The resin was polymerized by placing it at 60°C for 48 h. Ultrathin sections (80 nm) were prepared using an ultramicrotome (Ultracut UCT; Leica) and a diamond knife and embedded in copper grids. The sections were stained with 2% uranyl acetate at 25°C for 15 min, washed with distilled water, and stained with a lead staining solution (Sigma-Aldrich) at 25°C for 3 min. Sections were observed at an accelerating voltage of 100 kV using a TEM (JEM-1400 Plus; JEOL), and images were captured using a CCD camera (EM-14830RUBY2; JEOL).

### Transient expression and subcellular localization of orf320

The full-length cDNA of *orf320* and *MTP-orf320* were amplified from the *MTP-orf320* construct. cDNA was inserted into the pENTR/D-TOPO vector (Thermo Fisher Scientific) using an In-Fusion HD Cloning Kit (Takara Bio). The inserted sequences were cloned into a Ti plasmid using the Gateway LR reaction (Thermo Fisher Scientific) with the pUGW_sfGFP_HSPT vector.

Protoplasts were prepared from leaves of *Nicotiana benthamiana* and transformed as previously described (Lin et al. 2018). Four micrograms of each plasmid were used for polyethylene glycol (PEG)/glycol/calcium-mediated transformation. The protoplasts were treated with 200 nM MitoTracker Red CMXRos (Invitrogen) for 30 min. Protoplasts were observed under a confocal laser scanning microscope (LSM700; Zeiss). For the detection of the GFP signal, excitation was performed at 488 nm, whereas detection was performed between 490 and 555 nm. Similarly, to detect the MitoTracker Red signal, excitation was performed at 555 nm and detection was performed between 505 and 600 nm.

### PCR analysis for orf320 detection

Amplification was performed in 10 μL volume, containing 2 μL of total DNA (5 ng/μL), 5 μL of 2× Ampdirect Plus (Shimadzu), 0.25 U of BIOTAQ HS DNA Polymerase (Bioline Ltd.), and 0.3 μM forward and reverse primers (Supplementary Table S3). The thermal profile included initial incubation at 95°C for 10 min, followed by 35 cycles involving incubation at 95, 60, and 72°C for 30 s, 30 s, and 1 min, respectively, and a final extension at 72°C for 5 min.

## Data Availability

The mtDNA of the ‘May Queen’ cultivar was deposited in the DNA Data Bank of Japan (DDBJ) under the accession number LC867501-LC867503. The raw RNA-Seq sequences are available in the Sequence Read Archive (SRA) database of DDBJ under BioProject accession number PRJDB20298. The raw sequences of PacBio were deposited under the BioProject accession number PRJDB20319.

## Acknowledgments

We thank all the technical and administrative members of the T-PIRC Center at the University of Tsukuba and the Kazusa DNA Research Institute. We are also grateful to K. Tanaka at Obihiro University of Agriculture and Veterinary Medicine, for technical assistance. The seeds of the MicroTom cultivar (TOMJPF0001) were provided by the National BioResource Project Tomato. We thank Editage (www.editage.jp) for English language editing.

## Author Contributions

T.A. and K.K. conceived of and coordinated the study. R.N. and R.S. conducted the plant experiments. R.N., K.S., and K.K. performed sequencing analysis. R.N., R.S., K.S., and K.K. wrote the manuscript.

