## Supplemental Table for "Relationship Between Open Reading Frame 320, a Gene Causing Male Sterility in Tomatoes, and Cytoplasmic Male Sterility in Potatoes"

**Supplemental Table 1** Potato lines used for mitochondrial DNA comparative analysis

| Line name | Accession | cpDNA type | mtDNA type |
|---|---|---|---|
| *Solanum tuberosum* subsp. *andigenum* isolate ADG2 | MW122970-MW122972 | A | β** |
| *Solanum chaucha* isolate CHA | MW122976-MW122978 | A | β** |
| *Solanum juzepczukii* isolate JUZ | MW122973-MW122975 | C | α** |
| *Solanum ahanhuiri* isolate AJH | MW122955-MW122957 | C | β** |
| *Solanum tuberosum* subsp. *andigenum* isolate ADG1 | MW122967-MW122969 | C | β** |
| *Solanum stenotomum* subsp. *goniocalyx* isolate GON1 | MW122949-MW122951 | S | β** |
| *Solanum stenotomum* subsp. *goniocalyx* isolate GON2 | MW122952-MW122954 | S | β** |
| *Solanum phureja* isolate PHU | MW122958-MW122960 | S | β** |
| *Solanum stenotomum* subsp. *stenotomum* isolate STN | MW122961-MW122963 | S | β** |
| *Solanum bukasovii* isolate BUK1 | MW122964-MW122966 | S | β** |
| *Solanum curtilobum* isolate CUR | MW122979-MW122981 | S | β** |
| *Solanum tuberosum* clone 11379-03 | MW594267-MW594269 | S | β |
| *Solanum tuberosum* clone 12120-03 | MW594270-MW594272 | S | β |
| *Solanum tuberosum* clone 12625-02 | MW594273-MW594275 | S | β |
| *Solanum tuberosum* clone W5281.2 | MW594276-MW594278 | S | β |
| *Solanum tuberosum* cultivar PT56 | MF989953-MF989957 | T | β** |
| *Solanum tuberosum* cultivar Désirée | MN104801-MN104803 | T | β** |
| *Solanum tuberosum* clone 07506-01 | MW594251-MW594254 | T | β |
| *Solanum tuberosum* clone 08675-21 | MW594255-MW594258 | T | β |
| *Solanum tuberosum* clone H412-1 | MW594259-MW594262 | T | β |
| *Solanum tuberosum* clone DW84-1457 | MW594263-MW594266 | T | β |
| *Solanum tuberosum* cultivar May Queen | LC867501-LC867503 | T | β |
| *Solanum tuberosum* isolate TBR | MW122982-MW122984 | W | α** |
| *Solanum tuberosum* clone 10908-06 | MW594279-MW594281 | W | α |
| *Solanum tuberosum* clone 17H131 | LC649808-LC649812 | W* | α |
| *Solanum okadae* clone OKA15 | MW594282-MW594284 | W | γ |
| *Solanum tuberosum* Alwara | LC649829-LC649830 | W | γ |
| *Solanum tuberosum* 15H156 | LC649823-LC649825 | W* | γ |
| *Solanum tuberosum* 17H117-9 | LC649813-LC649819 | W | γ |
| *Solanum tuberosum* 18H225 | LC649820-LC649822 | W* | γ |

* The cytoplasmic type of these lines is D-type (Sanetomo et al. 2022), and the cp DNA type is W type according to Hosaka and Sanetomo 2012.

** Cytoplasmic type was determined by in silico analysis using a set of primers ALM_4/ALM_5 (L ̈ossl et al. 2000)

**Supplemental Table 2** PCR analysis for *orf320* in wild species and relatives including landraces

| *Species** | Accesssion | Country | Region | Latitude | Longitude | cpDNA type | *orf320* |
|---|---|---|---|---|---|---|---|
| *Series Tuberosa* | | | | | | | |
| *Solanum achacachense* | PI 558032 | Bolivia | | | | | - |
| *Solanum alandiae* | PI 243501 | Bolivia | Cochabamba | | | | - |
| | PI 498085 | Bolivia | Cochabamba | -18.10000000 | -65.11666667 | | - |
| | PI 498086 | Bolivia | Santa Cruz | -18.68333333 | -64.18333333 | | + |
| | PI 498087 | Bolivia | Cochabamba | -18.11666667 | -65.10000000 | | - |
| | PI 498088 | Bolivia | Cochabamba | -17.73333333 | -65.10000000 | | - |
| | PI 498089 | Bolivia | Cochabamba | -17.71666667 | -65.10000000 | | - |
| | PI 498090 | Bolivia | Cochabamba | -17.71666667 | -65.10000000 | | + |
| | PI 498205 | Bolivia | Cochabamba | | | | - |
| | PI 527889 | Bolivia | Chuquisaca | -19.75000000 | -64.55000000 | | + |
| | PI 545844 | Bolivia | Cochabamba | -17.90000000 | -65.10000000 | | - |
| | PI 545845 | Bolivia | Cochabamba | -18.15000000 | -65.15000000 | | - |
| | PI 568913 | Bolivia | Cochabamba | | | | - |
| | PI 597728 | Bolivia | Cochabamba | -17.70000000 | -65.18333333 | | + |
| | PI 597729 | Bolivia | Cochabamba | -17.70000000 | -65.18333333 | | + |
| | PI 597730 | Bolivia | Cochabamba | -17.91666667 | -65.15000000 | | - |
| *Solanum avilesii* | PI 498091 | Bolivia | Santa Cruz | -18.63333333 | -64.15000000 | | - |
| | PI 498092 | Bolivia | Santa Cruz | -18.63333333 | -64.15000000 | | - |
| | PI 498093 | Bolivia | Santa Cruz | -18.63333333 | -64.15000000 | | + |
| | PI 568968 | Bolivia | Santa Cruz | | | | - |
| *Solanum berthaultii* | PI 218215 | Bolivia | Potosí | -19.58333333 | -65.75000000 | | - |
| | PI 283069 | Bolivia | Chuquisaca | | | | - |
| | PI 283070 | Bolivia | Chuquisaca | | | | - |
| | PI 473333 | Bolivia | Chuquisaca | -19.10000000 | -65.23333333 | | - |
| | PI 473334 | Bolivia | Chuquisaca | -19.03333333 | -65.28333333 | | - |
| | PI 473335 | Bolivia | Chuquisaca | -19.16666667 | -65.28333333 | | - |
| | PI 473337 | Bolivia | Potosí | -19.53333333 | -65.26666667 | | + |
| | PI 473338 | Bolivia | Potosí | -19.33333333 | -65.16666667 | | - |
| | PI 473339 | Bolivia | Chuquisaca | -19.16666667 | -65.28333333 | | + |
| | PI 473340 | Bolivia | Chuquisaca | -19.03333333 | -65.28333333 | | - |
| | PI 498075 | Bolivia | Potosí | -19.56666667 | -65.45000000 | | + |
| | PI 498141 | Bolivia | Chuquisaca | -19.25000000 | -64.50000000 | | - |
| | PI 527884 | Bolivia | Chuquisaca | -18.85000000 | -65.23333333 | | - |
| | PI 527886 | Bolivia | Chuquisaca | -19.21666667 | -65.16666667 | | - |
| | PI 545850 | Bolivia | Chuquisaca | -19.26666667 | -65.18333333 | | - |
| | PI 545851 | Bolivia | Chuquisaca | -19.31666667 | -64.83333333 | | - |
| | PI 545852 | Bolivia | Chuquisaca | -19.20000000 | -64.66666667 | | + |
| | PI 545886 | Bolivia | Chuquisaca | -19.11666667 | -64.85000000 | | - |
| | PI 545890 | Bolivia | Potosí | -19.38333333 | -65.36666667 | | - |
| | PI 545962 | Bolivia | Chuquisaca | -19.30000000 | -64.85000000 | | - |
| | PI 208881 | Argentina | | | | | - |
| | PI 265857 | Bolivia | Cochabamba | -17.40000000 | -66.15000000 | | - |
| | PI 265858 | Bolivia | Cochabamba | -17.40000000 | -66.15000000 | | - |
| | PI 310926 | Bolivia | Cochabamba | -17.40000000 | -66.15000000 | | - |
| | PI 310927 | Bolivia | Cochabamba | -17.40000000 | -66.15000000 | | + |
| | PI 473330 | Bolivia | Cochabamba | -17.56666667 | -66.35000000 | | - |
| | PI 473331 | Bolivia | Cochabamba | -17.40000000 | -66.15000000 | | - |
| | PI 498094 | Bolivia | Cochabamba | -18.28333333 | -65.21666667 | | - |
| | PI 498109 | Bolivia | Cochabamba | -17.56666667 | -66.38333333 | | - |
| | PI 545849 | Bolivia | Cochabamba | -17.71666667 | -65.20000000 | | - |
| | PI 545885 | Bolivia | Cochabamba | -18.15000000 | -65.13333333 | | + |
| | PI 545960 | Bolivia | Cochabamba | -17.51666667 | -66.30000000 | | - |
| | PI 545961 | Bolivia | Cochabamba | -17.63333333 | -66.70000000 | | - |
| | PI 558033 | Bolivia | Cochabamba | -17.91666667 | -65.91666667 | | - |
| | PI 568918 | Bolivia | Cochabamba | | | | + |
| | PI 568920 | Bolivia | Santa Cruz | | | | - |
| *Solanum brevicaule* | PI 498111 | Bolivia | Cochabamba | -17.21666667 | -66.05000000 | W | + |
| | PI 498112 | Bolivia | Cochabamba | -17.21666667 | -66.05000000 | W | - |
| | PI 545967 | Bolivia | Cochabamba | -17.63333333 | -66.73333333 | W | - |
| | PI 545969 | Bolivia | Cochabamba | -17.61666667 | -66.71666667 | W | - |
| | PI 310929 | Bolivia | Cochabamba | -17.26666667 | -66.30000000 | W | - |
| | PI 310930 | Bolivia | Cochabamba | -17.26666667 | -66.30000000 | W | - |
| | PI 498110 | Bolivia | Cochabamba | -17.33333333 | -66.35000000 | W | - |
| | PI 545970 | Bolivia | La Paz | -15.60000000 | -69.01666667 | W | - |
| | PI 498218 | Bolivia | La Paz | -15.78333333 | -68.16666667 | W | + |
| | PI 498115 | Bolivia | Cochabamba | -17.33333333 | -66.35000000 | W | - |
| | PI 498114 | Bolivia | Cochabamba | -17.31666667 | -66.36666667 | W | - |
| | PI 545968 | Bolivia | Cochabamba | -17.63333333 | -66.65000000 | W | - |
| | PI 473378 | Bolivia | Cochabamba | -17.33333333 | -66.35000000 | W | - |
| | PI 498113 | Bolivia | Cochabamba | -17.21666667 | -66.05000000 | W | - |
| *Solanum candolleanum* | PI 568969 | Bolivia | La Paz | | | C | - |
| | PI 498227 | Bolivia | La Paz | -15.78333333 | -68.66666667 | C | - |
| *Solanum × doddsii* | PI 442690 | Bolivia | Cochabamba | -18.66666667 | -65.16666667 | | - |
| | PI 473350 | Bolivia | Cochabamba | | | | + |
| | PI 537028 | Bolivia | Cochabamba | -18.41666667 | -65.16666667 | | - |
| | PI 545854 | Bolivia | Cochabamba | -17.73333333 | -65.10000000 | | - |
| | PI 545855 | Bolivia | Cochabamba | -17.90000000 | -65.10000000 | | - |

| | | | | | | | |
|---|---|---|---|---|---|---|---|
| | PI 545856 | Bolivia | Cochabamba | -18.16666667 | -65.16666667 | | - |
| | PI 545857 | Bolivia | Cochabamba | -18.16666667 | -65.16666667 | | - |
| | PI 545858 | Bolivia | Cochabamba | -18.15000000 | -65.13333333 | | - |
| | PI 545859 | Bolivia | Cochabamba | -18.15000000 | -65.10000000 | | - |
| | PI 545860 | Bolivia | Cochabamba | -18.15000000 | -65.11666667 | | - |
| | PI 545861 | Bolivia | Cochabamba | -17.98333333 | -65.11666667 | | + |
| *Solanum gandarillasii* | PI 265866 | Bolivia | Cochabamba | -18.33333333 | -65.16666667 | | - |
| | PI 283076 | Bolivia | Cochabamba | -18.66666667 | -65.16666667 | | - |
| | PI 545862 | Bolivia | Cochabamba | -17.71666667 | -65.20000000 | | - |
| | PI 545863 | Bolivia | Cochabamba | -18.46666667 | -65.21666667 | | - |
| | PI 545864 | Bolivia | Cochabamba | -18.45000000 | -65.21666667 | | - |
| | PI 597750 | Bolivia | Chuquisaca | -18.91666667 | -65.10000000 | | - |
| | PI 597751 | Bolivia | Cochabamba | -18.50000000 | -65.16666667 | | - |
| *Solanum gourlayi* | | | | | | | |
| ssp. *gourlayi* | PI 473004 | Argentina | Jujuy | -23.56666667 | -65.43333333 | | - |
| | PI 473005 | Argentina | Jujuy | -23.56666667 | -65.43333333 | | - |
| | PI 473006 | Argentina | Jujuy | -23.56666667 | -65.43333333 | | - |
| | PI 473007 | Argentina | Jujuy | -23.56666667 | -65.43333333 | | - |
| ssp. *pachytrichum* | PI 545865 | Bolivia | Potosí | -19.56694444 | -65.38277778 | | + |
| | PI 545866 | Bolivia | Potosí | -19.56694444 | -65.38277778 | | - |
| | PI 545867 | Bolivia | Potosí | -19.56694444 | -65.36694444 | | + |
| | PI 545975 | Bolivia | Chuquisaca | -18.93333333 | -65.38333333 | | - |
| | PI 545976 | Bolivia | Chuquisaca | -18.93333333 | -65.38333333 | | - |
| | PI 545977 | Bolivia | Chuquisaca | -18.95000000 | -65.36666667 | | + |
| | PI 545978 | Bolivia | Chuquisaca | -18.95000000 | -65.31666667 | | + |
| ssp. *vidaurrei* | PI 472911 | Argentina | Salta | -22.20000000 | -65.16666667 | | + |
| | PI 472912 | Argentina | Salta | -22.18333333 | -65.18333333 | | + |
| | PI 472991 | Argentina | Jujuy | -23.20000000 | -65.45000000 | | + |
| | PI 472994 | Argentina | Salta | -22.71666667 | -65.25000000 | | + |
| | PI 472995 | Argentina | Salta | -22.71666667 | -65.20000000 | | + |
| | PI 472996 | Argentina | Salta | -22.68333333 | -65.20000000 | | + |
| | PI 472997 | Argentina | Salta | -22.66666667 | -65.20000000 | | + |
| | PI 472998 | Argentina | Salta | -22.60000000 | -65.16666667 | | + |
| | PI 472999 | Argentina | Salta | -22.51666667 | -65.10000000 | | - |
| | PI 473000 | Argentina | Salta | -22.51666667 | -65.10000000 | | - |
| | PI 473104 | Argentina | Salta | -22.26666667 | -65.20000000 | | - |
| | PI 473105 | Argentina | Salta | -22.18333333 | -65.18333333 | | - |
| | PI 473106 | Argentina | Salta | -22.18333333 | -65.18333333 | | + |
| | PI 473107 | Argentina | Salta | -22.18333333 | -65.18333333 | | - |
| | PI 498332 | Argentina | Salta | -22.81666667 | -65.23333333 | | + |
| | PI 597712 | Bolivia | Chuquisaca | -20.36666667 | -65.11666667 | | + |
| *Solanum hondelmannii* | PI 473365 | Bolivia | Chuquisaca | -19.16666667 | -65.28333333 | | - |
| | PI 473366 | Bolivia | Chuquisaca | -19.16666667 | -65.28333333 | | - |
| | PI 498071 | Bolivia | Potosí | -19.33333333 | -65.16666667 | | - |
| | PI 498281 | Bolivia | Chuquisaca | | | | - |
| | PI 545868 | Bolivia | Chuquisaca | -19.26666667 | -65.18333333 | | - |
| | PI 545869 | Bolivia | Chuquisaca | -19.26666667 | -65.18333333 | | - |
| | PI 545870 | Bolivia | Chuquisaca | -19.26666667 | -65.18333333 | | - |
| | PI 545871 | Bolivia | Chuquisaca | -19.23333333 | -65.16666667 | | - |
| | PI 545872 | Bolivia | Chuquisaca | -19.23333333 | -65.16666667 | | - |
| | PI 545873 | Bolivia | Chuquisaca | -19.23333333 | -65.16666667 | | - |
| | PI 545874 | Bolivia | Chuquisaca | -19.25000000 | -65.16666667 | | - |
| | PI 545875 | Bolivia | Chuquisaca | -19.36666667 | -64.78333333 | | - |
| | PI 545877 | Bolivia | Potosí | -19.41666667 | -65.40000000 | | - |
| | PI 545878 | Bolivia | Potosí | -19.41666667 | -65.40000000 | | - |
| | PI 545879 | Bolivia | Cochabamba | -17.98333333 | -65.11666667 | | - |
| *Solanum hoopesii* | PI 545881 | Bolivia | Chuquisaca | -19.55000000 | -64.65000000 | | + |
| | PI 545882 | Bolivia | Chuquisaca | -19.48333333 | -64.70000000 | | + |
| | PI 597721 | Bolivia | Chuquisaca | -19.88333333 | -64.56666667 | | + |
| | PI 597752 | Bolivia | Chuquisaca | -20.10000000 | -64.41666667 | | - |
| *Solanum incamayoense* | PI 473060 | Argentina | Salta | -24.81666667 | -65.70000000 | | - |
| | PI 473066 | Argentina | Salta | -24.76666667 | -65.73333333 | | - |
| | PI 473067 | Argentina | Salta | -24.75000000 | -65.73333333 | | - |
| | PI 473068 | Argentina | Salta | -24.71666667 | -65.75000000 | | - |
| | PI 473069 | Argentina | Salta | -24.66666667 | -65.76666667 | | - |
| | PI 473070 | Argentina | Salta | -24.51666667 | -65.85000000 | | - |
| | PI 473089 | Argentina | Salta | -24.78333333 | -65.71666667 | | - |
| | PI 500048 | Argentina | Salta | -24.85000000 | -65.71666667 | | - |
| *Solanum kurtzianum* | PI 472928 | Argentina | Catamarca | -27.40000000 | -66.93333333 | | + |
| | PI 472930 | Argentina | Catamarca | -27.56666667 | -66.98333333 | | - |
| | PI 472934 | Argentina | Catamarca | -27.46666667 | -66.43333333 | | - |
| | PI 472935 | Argentina | Catamarca | -27.46666667 | -66.43333333 | | - |
| | PI 472936 | Argentina | Catamarca | -27.46666667 | -66.43333333 | | - |
| | PI 472941 | Argentina | Catamarca | -27.65000000 | -66.18333333 | | + |
| | PI 472937 | Argentina | Catamarca | -27.46666667 | -66.43333333 | | + |
| | PI 472942 | Argentina | Catamarca | -27.61666667 | -66.16666667 | | + |
| | PI 472953 | Argentina | Catamarca | -27.90000000 | -67.36666667 | | + |
| | PI 472959 | Argentina | Catamarca | -27.95000000 | -67.20000000 | | + |
| *Solanum leptophyes* | PI 473451 | Peru | Ayacucho | -15.55000000 | -73.63333333 | C | - |

| | | | | | | | |
|---|---|---|---|---|---|---|---|
| | PI 545990 | Bolivia | Potosí | -18.01666667 | -66.36666667 | C | - |
| | PI 473445 | Peru | Cusco | -13.51666667 | -71.98333333 | C | - |
| | PI 545986 | Bolivia | Oruro | -17.96666667 | -66.93333333 | W | + |
| | PI 545989 | Bolivia | Potosí | -18.01666667 | -66.36666667 | W | - |
| | PI 545988 | Bolivia | Potosí | -18.01666667 | -66.36666667 | W | - |
| | PI 545985 | Bolivia | Oruro | -17.96666667 | -66.93333333 | W | + |
| | PI 320340 | Bolivia | Oruro | | | W | + |
| | PI 545984 | Bolivia | Oruro | -17.96666667 | -66.93333333 | W | + |
| | PI 545991 | Bolivia | Potosí | -18.01666667 | -66.36666667 | W | - |
| | PI 545992 | Bolivia | Potosí | -18.01666667 | -66.36666667 | W | + |
| | PI 545993 | Bolivia | Potosí | -18.01666667 | -66.36666667 | W | + |
| | PI 545994 | Bolivia | Potosí | -18.01666667 | -66.36666667 | W | + |
| | PI 458378 | Peru | Puno | -16.36954700 | -69.22142500 | W | - |
| | PI 545995 | Bolivia | Potosí | -18.01666667 | -66.38333333 | W | - |
| | PI 545996 | Bolivia | Potosí | -18.01666667 | -66.38333333 | W | - |
| | PI 473343 | Bolivia | La Paz | -16.58333333 | -68.11666667 | W | - |
| | PI 473342 | Bolivia | La Paz | -16.55000000 | -68.10000000 | W | + |
| | PI 473344 | Bolivia | La Paz | -17.26666667 | -68.71666667 | W | - |
| | PI 283090 | Bolivia | La Paz | -16.30000000 | -68.13333300 | W | - |
| | PI 545987 | Bolivia | Potosí | -18.01666667 | -66.36666667 | W | - |
| | PI 545895 | Bolivia | Potosí | -19.56694444 | -65.38277778 | W2 | - |
| | PI 545896 | Bolivia | Potosí | -19.56694444 | -65.36694444 | W2 | + |
| *Solanum microdontum* | PI 320307 | Argentina | Salta | -22.25000000 | -65.03333333 | | + |
| | PI 320309 | Argentina | Salta | -22.25000000 | -65.03333333 | | + |
| | PI 320310 | Argentina | Salta | -22.25000000 | -65.05000000 | | + |
| | PI 320314 | Argentina | Salta | -25.70000000 | -65.50000000 | | + |
| | PI 320316 | Argentina | Salta | -24.75000000 | -65.46666667 | | + |
| | PI 458356 | Argentina | Salta | -23.21666667 | -64.91666667 | | + |
| | PI 458357 | Argentina | Salta | -23.21666667 | -64.91666667 | | - |
| | PI 458358 | Argentina | Salta | -25.15000000 | -65.75000000 | | + |
| | PI 473166 | Argentina | Salta | -22.18333333 | -65.05000000 | | + |
| | PI 473167 | Argentina | Salta | -23.21666667 | -64.91666667 | | + |
| | PI 473168 | Argentina | Salta | -25.15000000 | -65.65000000 | | + |
| | PI 473169 | Argentina | Salta | -22.25000000 | -65.03333333 | | + |
| | PI 473170 | Argentina | Salta | -24.90000000 | -65.65000000 | | + |
| | PI 473171 | Argentina | Salta | -25.15000000 | -65.68333333 | | + |
| | PI 473172 | Argentina | Salta | -22.20000000 | -65.03333333 | | + |
| | PI 473173 | Argentina | Salta | -22.15000000 | -65.03333333 | | + |
| | PI 473174 | Argentina | Salta | -22.13333333 | -65.03333333 | | - |
| | PI 473175 | Argentina | Salta | -22.13333333 | -65.03333333 | | - |
| | PI 473176 | Argentina | Salta | -23.11694444 | -64.91694444 | | - |
| | PI 473177 | Argentina | Salta | -22.20000000 | -65.03333333 | | + |
| | PI 473179 | Argentina | Salta | -22.33333333 | -65.03333333 | | - |
| | PI 473180 | Argentina | Salta | -22.33333333 | -65.03333333 | | - |
| | PI 473312 | Argentina | Salta | -22.73333333 | -65.23333333 | | + |
| | PI 500032 | Argentina | Salta | -25.18333333 | -65.80000000 | | + |
| | PI 500033 | Argentina | Salta | -24.63333333 | -65.50000000 | | + |
| | PI 500034 | Argentina | Salta | -24.61666667 | -65.56666667 | | - |
| | PI 500035 | Argentina | Salta | -24.61666667 | -65.51666667 | | + |
| | PI 500036 | Argentina | Salta | -24.61666667 | -65.58333333 | | - |
| | PI 500037 | Argentina | Salta | -24.63333333 | -65.48333333 | | + |
| | PI 500038 | Argentina | Salta | -25.18333333 | -65.80000000 | | + |
| | PI 500041 | Argentina | Salta | -25.18333333 | -65.85000000 | | + |
| | PI 500044 | Argentina | Salta | -24.61666667 | -65.58333333 | | - |
| | PI 500064 | Argentina | Salta | -24.61666667 | -65.56666667 | | + |
| | PI 558097 | Argentina | Salta | -24.61666667 | -65.58333333 | | + |
| | PI 558098 | Argentina | Salta | -24.60000000 | -65.58333333 | | + |
| | PI 595508 | Argentina | Salta | | | | + |
| | PI 320318 | Argentina | Salta | -25.15000000 | -65.86666667 | | + |
| | PI 473362 | Bolivia | Santa Cruz | -18.71353300 | -64.14881700 | | - |
| | PI 498121 | Bolivia | Chuquisaca | -19.41666667 | -64.26666667 | | + |
| | PI 498123 | Bolivia | Chuquisaca | -19.01666667 | -64.31666667 | | - |
| | PI 498124 | Bolivia | Santa Cruz | -18.58277778 | -64.23277778 | | - |
| | PI 498125 | Bolivia | Santa Cruz | -18.58277778 | -64.23277778 | | - |
| | PI 498126 | Bolivia | Santa Cruz | -18.60000000 | -64.11666667 | | - |
| | PI 498127 | Bolivia | Santa Cruz | -18.61666667 | -64.13333333 | | - |
| | PI 498128 | Bolivia | Santa Cruz | -17.88333333 | -64.70000000 | | - |
| | PI 545901 | Bolivia | Tarija | -21.48333333 | -64.90000000 | | - |
| | PI 545904 | Bolivia | Chuquisaca | -19.75000000 | -64.55000000 | | + |
| | PI 595505 | Bolivia | | | | | + |
| | PI 595510 | Bolivia | Tarija | -21.40000000 | -64.46666667 | | + |
| | PI 595511 | Bolivia | Tarija | -21.40000000 | -64.43333333 | | + |
| | PI 597756 | Bolivia | Tarija | -21.45000000 | -64.35000000 | | + |
| | PI 597757 | Bolivia | Tarija | -21.40000000 | -64.30000000 | | + |
| *Solanum neocardenasii* | PI 498129 | Bolivia | Santa Cruz | -18.11666667 | -64.20000000 | | - |
| | PI 502642 | Bolivia | Santa Cruz | -18.10000000 | -64.18333333 | | - |
| *Solanum neorossii* | PI 473201 | Argentina | Salta | -22.25000000 | -65.03333333 | | - |
| | PI 473202 | Argentina | Salta | -22.25000000 | -65.08333333 | | + |
| | PI 473428 | Argentina | Salta | -22.25000000 | -65.06666667 | | + |
| | PI 473429 | Argentina | Salta | -22.25000000 | -65.06666667 | | + |

| | | | | | | | |
|---|---|---|---|---|---|---|---|
| | PI 473529 | Argentina | | | | | + |
| *Solanum okadae* | PI 320327 | Argentina | Salta | -25.15000000 | -65.86666667 | | - |
| | PI 320328 | Argentina | Salta | -25.15000000 | -65.86666667 | | - |
| | PI 458367 | Argentina | Salta | -25.15000000 | -65.83333333 | | - |
| | PI 458368 | Argentina | Salta | -25.15000000 | -65.83333333 | | - |
| | PI 498063 | Bolivia | La Paz | -17.03333333 | -67.25000000 | | - |
| | PI 498064 | Bolivia | La Paz | -17.03333333 | -67.25000000 | | - |
| | PI 498065 | Bolivia | La Paz | -17.03333333 | -67.25000000 | | - |
| | PI 498130 | Bolivia | Cochabamba | -17.10000000 | -66.91666667 | | + |
| | PI 498403 | Argentina | Jujuy | -23.63333333 | -65.10000000 | | - |
| | PI 498404 | Argentina | Jujuy | -23.63333333 | -65.10000000 | | - |
| | PI 500061 | Argentina | Salta | -25.18333333 | -65.85000000 | | - |
| | PI 558102 | Argentina | Salta | -25.15000000 | -65.85000000 | | - |
| | PI 558103 | Argentina | Salta | -24.61666667 | -65.56666667 | | - |
| | PI 558104 | Argentina | Jujuy | -24.11666667 | -65.63333333 | | - |
| | PI 558105 | Argentina | Jujuy | -23.60000000 | -65.18333333 | | - |
| | PI 597759 | Bolivia | Cochabamba | -16.95000000 | -67.18333333 | | - |
| *Solanum oplocense* | PI 435080 | Argentina | Jujuy | -21.88333333 | -66.18333333 | | + |
| | PI 442682 | Argentina | Jujuy | -21.88333333 | -66.18333333 | | + |
| | PI 442683 | Argentina | Jujuy | -21.88333333 | -66.18333333 | | + |
| | PI 442684 | Argentina | Jujuy | -21.88333333 | -66.18333333 | | + |
| | PI 442685 | Argentina | Jujuy | -21.88333333 | -66.18333333 | | + |
| | PI 442693 | Bolivia | Potosí | -19.50000000 | -65.81666667 | | + |
| | PI 458359 | Argentina | Jujuy | -22.11666667 | -65.46666667 | | + |
| | PI 458361 | Argentina | Jujuy | -21.88333333 | -66.18333333 | | + |
| | PI 458363 | Argentina | Jujuy | -21.88333333 | -66.18333333 | | + |
| | PI 458390 | Bolivia | Chuquisaca | -20.63333333 | -65.23333333 | W | - |
| | PI 473182 | Argentina | Jujuy | -21.88333333 | -66.18333333 | | + |
| | PI 473183 | Argentina | Jujuy | -21.88333333 | -66.18333333 | | + |
| | PI 473184 | Argentina | Jujuy | -21.88333333 | -66.18333333 | | + |
| | PI 473185 | Argentina | Jujuy | -21.88333333 | -66.18333333 | | + |
| | PI 473189 | Argentina | Jujuy | -22.10000000 | -65.46666667 | | + |
| | PI 473190 | Argentina | Jujuy | -22.10000000 | -65.46666667 | | + |
| | PI 473191 | Argentina | Jujuy | -22.10000000 | -65.46666667 | | + |
| | PI 473192 | Argentina | Jujuy | -21.88333333 | -66.18333333 | | + |
| | PI 473193 | Argentina | Jujuy | -21.88333333 | -66.18333333 | | + |
| | PI 473194 | Argentina | Jujuy | -21.88333333 | -66.18333333 | | + |
| | PI 473195 | Argentina | Jujuy | -21.88333333 | -66.18333333 | | + |
| | PI 473198 | Argentina | Jujuy | -22.11666667 | -65.46666667 | | + |
| | PI 473199 | Argentina | Jujuy | -22.11666667 | -65.46666667 | | + |
| | PI 473499 | Bolivia | Potosí | -20.93333333 | -66.35000000 | | + |
| | PI 473500 | Bolivia | Potosí | -20.93333333 | -66.35000000 | | + |
| | PI 498068 | Bolivia | Potosí | -21.71666667 | -65.65000000 | | + |
| | PI 498069 | Bolivia | Potosí | -20.93333333 | -66.35000000 | | + |
| | PI 498070 | Bolivia | Potosí | -20.83333333 | -65.68333333 | | + |
| | PI 498269 | Bolivia | Potosí | -21.33333333 | -65.66666667 | | + |
| | PI 498270 | Bolivia | Potosí | -21.50000000 | -65.58333333 | | + |
| | PI 498389 | Argentina | Jujuy | -21.88333333 | -66.18333333 | | + |
| | PI 545906 | Bolivia | Potosí | -21.38333333 | -65.81666667 | | + |
| | PI 545907 | Bolivia | Potosí | -21.36666667 | -65.85000000 | | + |
| | PI 545909 | Bolivia | Potosí | -21.13333333 | -65.80000000 | | + |
| *Solanum oplocense* | PI 545876 | Bolivia | Potosí | -19.43333333 | -65.40000000 | W | - |
| | PI 435079 | Argentina | Jujuy | -23.20000000 | -65.45000000 | W | + |
| | PI 545910 | Bolivia | Cochabamba | -17.98333333 | -65.11666667 | W | + |
| | PI 545908 | Bolivia | Potosí | -21.13333333 | -65.80000000 | W | + |
| *Solanum oplocense* | PI 498067 | Bolivia | Chuquisaca | -19.10000000 | -65.23333333 | W2 | - |
| *Solanum sparsipilum* | | | | | | | |
| ssp. *sparsipilum* | PI 498136 | Bolivia | Cochabamba | -17.38333333 | -66.16666667 | W | - |
| | PI 498137 | Bolivia | Cochabamba | -18.11666667 | -65.10000000 | W | + |
| | PI 498140 | Bolivia | Cochabamba | -17.63333333 | -66.30000000 | W | + |
| | PI 498305 | Peru | Cusco | -13.33333333 | -71.90000000 | W | - |
| | PI 498282 | Bolivia | La Paz | -16.16666667 | -69.08333333 | W | - |
| | PI 498283 | Bolivia | La Paz | | | W | - |
| | PI 498138 | Bolivia | Cochabamba | -17.56666667 | -66.38333333 | W | + |
| | PI 498139 | Bolivia | Cochabamba | -17.61666667 | -66.31666667 | W | - |
| *Solanum spegazzinii* | PI 442686 | Argentina | Salta | -24.80000000 | -66.16666667 | | + |
| | PI 458335 | Argentina | Salta | -25.41666667 | -65.93333333 | | + |
| | PI 472967 | Argentina | Salta | -24.80000000 | -66.16666667 | | + |
| | PI 472968 | Argentina | Salta | -24.80000000 | -66.16666667 | | + |
| | PI 472977 | Argentina | Salta | -24.86666667 | -66.03333333 | | - |
| | PI 472980 | Argentina | Salta | -25.13333333 | -66.61666667 | | + |
| | PI 472969 | Argentina | Salta | -24.80000000 | -66.16666667 | | + |
| | PI 500050 | Argentina | Salta | -25.06666667 | -66.03333333 | | + |
| *Solanum × sucrense* | PI 258906 | Bolivia | Potosí | -19.76666667 | -65.46666667 | | - |
| | PI 290959 | Bolivia | Chuquisaca | -19.03333333 | -65.28333333 | | - |
| | PI 442691 | Bolivia | Potosí | -19.50000000 | -65.65000000 | | - |
| | PI 442692 | Bolivia | Potosí | -19.56666667 | -65.43333333 | | + |
| | PI 458391 | Bolivia | Potosí | -21.31666667 | -65.76666667 | | - |
| | PI 473364 | Bolivia | Potosí | -19.50000000 | -65.81666667 | | - |
| | PI 473367 | Bolivia | Chuquisaca | -20.50000000 | -65.16666667 | | - |

| | | | | | | | |
|---|---|---|---|---|---|---|---|
| | PI 473374 | Bolivia | Potosí | -19.63333333 | -65.06666667 | | - |
| | PI 473379 | Bolivia | Potosí | -19.63333333 | -65.06666667 | | + |
| | PI 473381 | Bolivia | Potosí | -19.56666667 | -65.43333333 | | + |
| | PI 473382 | Bolivia | Potosí | -19.56666667 | -65.43333333 | | + |
| | PI 473388 | Bolivia | Potosí | -19.56666667 | -65.43333333 | | + |
| | PI 473392 | Bolivia | Potosí | -19.63333333 | -65.06666667 | | + |
| | PI 473506 | Bolivia | Chuquisaca | -20.83277778 | -65.75000000 | | - |
| | PI 498286 | Bolivia | Chuquisaca | | | | + |
| | PI 498300 | Bolivia | Potosí | -19.46666667 | -65.83333333 | | + |
| | PI 498301 | Bolivia | Potosí | -19.46666667 | -65.83333333 | | + |
| | PI 498306 | Bolivia | Chuquisaca | -20.25000000 | -65.18333333 | | - |
| | PI 545887 | Bolivia | Chuquisaca | -20.23333333 | -65.11666667 | | + |
| | PI 545888 | Bolivia | Potosí | -21.13333333 | -65.80000000 | | - |
| | PI 545915 | Bolivia | Chuquisaca | -19.25000000 | -64.88333333 | | - |
| | PI 545918 | Bolivia | Potosí | -19.48333333 | -65.63333333 | | + |
| | PI 545929 | Bolivia | Chuquisaca | -19.25000000 | -65.16666667 | | - |
| | PI 546008 | Bolivia | Chuquisaca | -19.25000000 | -64.88333333 | | - |
| | PI 595512 | Bolivia | Potosí | -19.55000000 | -65.70000000 | | + |
| *Solanum ugentii* | PI 546029 | Bolivia | Chuquisaca | -19.60000000 | -64.61666667 | | + |
| | PI 546030 | Bolivia | Chuquisaca | -19.55000000 | -64.65000000 | | - |
| | PI 546032 | Bolivia | Chuquisaca | -19.51666667 | -64.68333333 | | + |
| *Solanum venturii* | PI 218220 | Argentina | Tucumán | | | | - |
| | PI 558146 | Argentina | Jujuy | -24.11666667 | -65.63333333 | | - |
| | PI 558224 | Argentina | Catamarca | -27.33333333 | -66.01666667 | | + |
| | PI 566792 | Argentina | Jujuy | -23.63333333 | -65.10000000 | | - |
| *Solanum vernei* | | | | | | | |
| ssp. *vernei* | PI 230468 | Argentina | Tucumán | | | | - |
| | PI 230562 | Argentina | Tucumán | | | | - |
| | PI 320330 | Argentina | Tucumán | -26.76666667 | -65.75000000 | | - |
| | PI 320332 | Argentina | Catamarca | -27.35000000 | -66.03333333 | | - |
| | PI 458373 | Argentina | Tucumán | -26.76666667 | -65.76666667 | | - |
| | PI 458374 | Argentina | Salta | -22.20000000 | -65.16666667 | W | - |
| | PI 473306 | Argentina | Salta | -25.16666667 | -65.86666667 | | + |
| | PI 473310 | Argentina | Salta | -22.11666667 | -65.03333333 | | - |
| | PI 473311 | Argentina | Salta | -22.03333333 | -65.03333333 | | - |
| | PI 500062 | Argentina | Salta | -25.15000000 | -65.85000000 | | - |
| | PI 500065 | Argentina | Salta | -24.60000000 | -65.60000000 | | - |
| | PI 500068 | Argentina | Jujuy | -23.58333333 | -65.21666667 | | - |
| | PI 500069 | Argentina | Jujuy | -23.60000000 | -65.18333333 | | - |
| | PI 558147 | Argentina | Salta | -25.15000000 | -65.85000000 | | - |
| | PI 558149 | Argentina | Jujuy | -24.06666667 | -65.65000000 | | - |
| | PI 558150 | Argentina | Jujuy | -23.60000000 | -65.18333333 | | - |
| | PI 500067 | Argentina | Jujuy | -24.05000000 | -65.60000000 | W | - |
| | PI 558151 | Argentina | Jujuy | -23.60000000 | -65.18333333 | W | - |
| | PI 473309 | Argentina | Salta | -23.11666667 | -64.53333333 | W | - |
| | PI 558148 | Argentina | Salta | -25.16666667 | -65.86666667 | W | - |
| | PI 545884 | Bolivia | Cochabamba | -17.66666667 | -65.30000000 | W | - |
| ssp. *ballsii* | PI 320333 | Argentina | Jujuy | -23.91666667 | -65.41666667 | | - |
| | PI 458369 | Argentina | Jujuy | -23.60000000 | -65.13333333 | | - |
| | PI 458370 | Argentina | Salta | -23.16666667 | -64.95000000 | | - |
| | PI 458371 | Argentina | Salta | -23.16666667 | -64.95000000 | | - |
| | PI 458372 | Argentina | Salta | -23.16666667 | -64.95000000 | | - |
| | PI 473303 | Argentina | Jujuy | -23.60000000 | -65.13333333 | | - |
| | PI 473304 | Argentina | Jujuy | -23.60000000 | -65.13333333 | | - |
| | PI 473305 | Argentina | Jujuy | -23.63333333 | -65.10000000 | | - |
| | PI 500070 | Argentina | Salta | -23.20000000 | -64.91666667 | | - |
| *Solanum virgultorum* | BGRC27159 | | | | | | - |
| | BGRC31203 | | | | | | - |
| | BGRC8265 | | | | | | - |
| | CGN20615 | | | | | | - |
| | CGN20652 | | | | | | - |
| *Series Megistacroloba* | | | | | | | |
| *Solanum astleyi* | PI 545848 | Bolivia | Potosí | -19.43333333 | -65.40000000 | | - |
| | PI 545959 | Bolivia | Potosí | -19.43333333 | -65.40000000 | | - |
| *Solanum boliviense* | PI 545964 | Bolivia | Potosí | -19.43333333 | -65.40000000 | W | + |
| | PI 498215 | Bolivia | Chuquisaca | -18.98333333 | -65.36666667 | W2 | + |
| | PI 265860 | Bolivia | Chuquisaca | -19.13333333 | -64.90000000 | | + |
| | PI 265861 | Bolivia | Chuquisaca | -19.13333333 | -64.90000000 | | - |
| | PI 310928 | Bolivia | Chuquisaca | -19.03333333 | -65.28333333 | | + |
| | PI 545889 | Bolivia | Potosí | -19.43333333 | -65.40000000 | | + |
| | PI 545963 | Bolivia | Potosí | -19.43333333 | -65.40000000 | | + |
| | PI 545965 | Bolivia | Potosí | -19.48333333 | -65.45000000 | | + |
| | PI 545979 | Bolivia | Potosí | -19.43333333 | -65.40000000 | | + |
| | PI 568921 | Bolivia | Chuquisaca | | | | + |
| *Solanum megistacrolobum* | PI 275148 | Argentina | Salta | -22.21666667 | -65.23333333 | | - |
| | PI 435077 | Argentina | Salta | -25.13333333 | -65.86666667 | | - |
| | PI 473158 | Argentina | Salta | -22.26666667 | -65.18333333 | | - |
| | PI 500029 | Argentina | Salta | -25.16666667 | -65.86666667 | | - |
| | PI 500030 | Argentina | Salta | -25.33333333 | -65.88333333 | | - |
| | PI 265874 | Bolivia | Potosí | -19.55000000 | -65.75000000 | C | - |

| | | | | | | | |
|---|---|---|---|---|---|---|---|
| | PI 473361 | Peru | Puno | -16.36954700 | -69.22142500 | C | - |
| | PI 473356 | Bolivia | Potosí | -19.63333333 | -65.16666667 | C | - |
| *Solanum sanctae-rosae* | PI 205397 | Argentina | | | | | - |
| | PI 218221 | Argentina | Tucumán | | | | - |
| | PI 230464 | Argentina | Tucumán | -26.78277778 | -65.75000000 | | - |
| | PI 275152 | Argentina | Tucumán | -26.78277778 | -65.75000000 | | - |
| | PI 283089 | Argentina | Tucumán | -26.55000000 | -65.80000000 | | - |
| | PI 320324 | Argentina | Tucumán | -26.73333333 | -65.73333333 | | - |
| | PI 320325 | Argentina | Salta | -25.83333333 | -65.58333333 | | - |
| | PI 473200 | Argentina | Tucumán | -26.76666667 | -65.75000000 | | + |
| | PI 498391 | Argentina | Tucumán | -26.76666667 | -65.75000000 | | - |
| *Solanum sogarandinum* | PI 230510 | Peru | La Libertad | -8.15000000 | -78.18333333 | | - |
| | PI 365360 | Peru | Ancash | | | | - |
| *Solanum toralapanum* | PI 195210 | Bolivia | Cochabamba | -17.43333333 | -65.71666667 | | - |
| | PI 310934 | Bolivia | Cochabamba | -18.50000000 | -65.28333333 | | - |
| | PI 320302 | Argentina | Salta | -25.28333333 | -65.90000000 | | - |
| | PI 320303 | Argentina | Salta | -22.25000000 | -65.08333333 | | - |
| | PI 458396 | Bolivia | Chuquisaca | -19.15000000 | -64.91666667 | | - |
| | PI 458397 | Bolivia | Tarija | -21.51666667 | -64.90000000 | | - |
| | PI 472804 | Argentina | Salta | -22.30000000 | -65.05000000 | | - |
| | PI 472805 | Argentina | Salta | -22.25000000 | -65.30000000 | | - |
| | PI 472806 | Argentina | Salta | -22.28333333 | -64.86666667 | | - |
| | PI 472807 | Argentina | Salta | -22.33333333 | -65.00000000 | | - |
| | PI 472808 | Argentina | Salta | -22.21666667 | -65.23333333 | | - |
| | PI 473389 | Bolivia | Cochabamba | -17.43333333 | -65.71666667 | | - |
| | PI 498142 | Bolivia | Cochabamba | -17.25000000 | -66.90000000 | | - |
| | PI 498143 | Bolivia | Cochabamba | -17.23333333 | -66.90000000 | | - |
| | PI 498144 | Bolivia | Cochabamba | -17.23333333 | -66.05000000 | | - |
| | PI 498145 | Bolivia | Cochabamba | -17.23333333 | -66.05000000 | | - |
| | PI 498146 | Bolivia | Cochabamba | -17.21666667 | -66.05000000 | | - |
| | PI 498257 | Bolivia | La Paz | -17.06666667 | -67.30000000 | | - |
| | PI 498263 | Bolivia | Cochabamba | | | | - |
| | PI 545892 | Bolivia | Cochabamba | -17.30000000 | -65.73333333 | | - |
| | PI 545925 | Bolivia | Cochabamba | -17.31666667 | -66.86666667 | | - |
| | PI 545926 | Bolivia | Tarija | -21.48333333 | -64.88333333 | | - |
| | PI 545927 | Bolivia | Cochabamba | -17.43333333 | -65.46666667 | | - |
| | PI 545928 | Bolivia | Cochabamba | -17.30000000 | -65.73333333 | | - |
| | PI 545998 | Bolivia | Potosí | -18.00000000 | -66.43333333 | | - |
| | PI 546009 | Bolivia | Cochabamba | -17.63333333 | -66.65000000 | | - |
| | PI 546010 | Bolivia | Cochabamba | -17.31666667 | -66.86666667 | | - |
| | PI 546011 | Bolivia | Cochabamba | -17.63333333 | -66.73333333 | | - |
| | PI 546012 | Bolivia | Cochabamba | -17.63333333 | -66.65000000 | | - |
| | PI 546013 | Bolivia | Cochabamba | -17.63333333 | -66.70000000 | | - |
| | PI 546014 | Bolivia | La Paz | -17.05000000 | -67.88333333 | | - |
| | PI 546015 | Bolivia | Chuquisaca | -19.60000000 | -64.61666667 | | - |
| | PI 607882 | Peru | Puno | -15.28111111 | -70.28138889 | | - |
| *Series Yungasensa* | | | | | | | |
| *Solanum arnezii* | PI 545846 | Bolivia | Chuquisaca | -19.60000000 | -64.65000000 | | - |
| | PI 545847 | Bolivia | Chuquisaca | -19.55000000 | -64.65000000 | | + |
| | PI 545880 | Bolivia | Chuquisaca | -19.60000000 | -64.63333333 | | - |
| | PI 545958 | Bolivia | Chuquisaca | -19.56666667 | -64.53333333 | | - |
| *Solanum chacoense* | PI 537025 | Bolivia | Chuquisaca | | | | + |
| *Solanum tarijense* | PI 217458 | Argentina | Salta | -23.00000000 | -64.56666667 | | - |
| | PI 275154 | Argentina | Salta | -22.25000000 | -64.96666667 | | - |
| | PI 414148 | Argentina | Salta | -22.21666667 | -64.95000000 | | - |
| | PI 414152 | Bolivia | Tarija | -21.51694444 | -64.55000000 | | - |
| | PI 442689 | Argentina | Salta | -22.21666667 | -64.95000000 | | - |
| | PI 458365 | Argentina | Salta | -22.25000000 | -64.88333333 | | - |
| | PI 458394 | Bolivia | Chuquisaca | -19.16666667 | -65.28333333 | | - |
| | PI 458395 | Bolivia | Tarija | -21.51694444 | -64.55000000 | | - |
| | PI 473217 | Argentina | Salta | -22.60000000 | -65.11666667 | | + |
| | PI 473227 | Argentina | Salta | -22.26666667 | -64.96666667 | | + |
| | PI 473332 | Bolivia | Chuquisaca | -19.10000000 | -65.23333333 | | - |
| | PI 473336 | Bolivia | Potosí | -19.33333333 | -65.16666667 | | + |
| | PI 498290 | Bolivia | Tarija | | | | - |
| | PI 498399 | Argentina | Salta | -22.25000000 | -64.96666667 | | + |
| | PI 500054 | Argentina | Salta | -25.16666667 | -65.81666667 | | - |
| | PI 545922 | Bolivia | Chuquisaca | -19.23333333 | -65.16666667 | | + |
| | PI 545923 | Bolivia | Chuquisaca | -19.25000000 | -65.16666667 | | - |
| | PI 558130 | Argentina | Salta | -25.18333333 | -65.80000000 | | - |
| | PI 597774 | Bolivia | Chuquisaca | -19.08333333 | -65.18333333 | | - |
| | PI 195206 | Bolivia | Chuquisaca | -19.81666667 | -63.98333333 | | - |
| | PI 217457 | Argentina | Salta | -22.21666667 | -64.88333333 | | + |
| | PI 265577 | Bolivia | Cochabamba | -17.93333333 | -65.16666667 | | - |
| | PI 414149 | Argentina | Salta | -22.20000000 | -64.95000000 | | - |
| | PI 414150 | Argentina | Salta | -22.25000000 | -64.96666667 | | + |
| | PI 458364 | Argentina | Salta | -22.25000000 | -64.96666667 | | + |
| | PI 458366 | Argentina | Salta | -22.25000000 | -64.95000000 | | - |
| | PI 472815 | Argentina | Salta | -25.18333333 | -65.80000000 | | - |
| | PI 473216 | Argentina | Salta | -22.65000000 | -65.18333333 | | - |

| | | | | | | | |
|---|---|---|---|---|---|---|---|
| | PI 473218 | Argentina | Salta | -22.25000000 | -64.96666667 | | - |
| | PI 473226 | Argentina | Salta | -22.26666667 | -64.96666667 | | - |
| | PI 473228 | Argentina | Salta | -22.26666667 | -64.96666667 | | - |
| | PI 473245 | Argentina | Salta | -22.26666667 | -64.98333333 | | - |
| | PI 500043 | Argentina | Salta | -25.18333333 | -65.78333333 | | - |
| | PI 500055 | Argentina | Salta | -25.18333333 | -65.78333333 | | - |
| | PI 545920 | Bolivia | Chuquisaca | -19.20000000 | -65.16666667 | | + |
| | PI 545921 | Bolivia | Chuquisaca | -19.23333333 | -65.18333333 | | - |
| | PI 545924 | Bolivia | Chuquisaca | -19.20000000 | -64.66666667 | | - |
| | PI 558129 | Argentina | Salta | -25.18333333 | -65.80000000 | | - |
| | PI 566799 | Argentina | Salta | | | | + |
| | PI 473232 | Argentina | Salta | -22.26666667 | -64.96666667 | T | - |
| | PI 473239 | Argentina | Salta | -22.26666667 | -64.96666667 | T | - |
| | PI 473243 | Argentina | Salta | -22.35000000 | -65.18333333 | T | + |
| *Solanum raphaniforium Cárd. et Hawkes (rap)* | PI 210048 | Peru | Cusco | -13.51666667 | -71.98333333 | C | - |
| | PI 473371 | Peru | Apurímac | -14.05000000 | -72.46666667 | C | - |
| *Solanum acaule Bitt. (acl)* | PI 210030 | Bolivia | Potosí | -19.63333333 | -65.06666667 | C | - |
| *Solanum acroglossum Juz. (acg)* | PI 498204 | Peru | Pasco | -10.66666667 | -76.08333333 | W | - |
| *Solanum blanco-galdosii Ochoa (blg)* | PI 442701 | Peru | Cajamarca | -9.93333333 | -78.21666667 | W | - |
| *Solanum acroscopicum Ochoa (acs)* | PI 365315 | Peru | Arequipa | -15.20000000 | -72.93333333 | C | - |
| | PI 365314 | Peru | Arequipa | -15.20000000 | -72.93333333 | W | - |
| *Solanum ambosinum Ochoa (amb)* | PI 365316 | Peru | Huánuco | -10.21666667 | -76.13333333 | C | - |
| *Solanum bukasovii Juz. (buk)* | PI 498222 | Peru | Junín | -11.80000000 | -75.50000000 | A | - |
| | PI 568944 | Peru | Pasco | | | C | - |
| | PI 568954 | Peru | Puno | | | A | - |
| | PI 473450 | Peru | Ayacucho | -15.25000000 | -73.71666667 | C | - |
| | PI 283074 | Peru | Cusco | -14.23333333 | -72.03333333 | C | - |
| | PI 230511 | Peru | Puno | -15.83333333 | -70.03333333 | C | - |
| | PI 498221 | Peru | Junín | -12.06666667 | -75.23333333 | C | - |
| | PI 365350 | Peru | Lima | -12.15000000 | -76.23333333 | C | - |
| | PI 568932 | Peru | Ayacucho | | | C | - |
| | PI 365349 | Peru | Lima | -12.36305556 | -75.85111111 | C | - |
| | PI 365304 | Peru | Lima | -12.15000000 | -76.23333333 | C | - |
| | PI 365321 | Peru | Huánuco | | | C | - |
| | PI 365355 | Peru | Cusco | -13.30000000 | -72.11666667 | C | - |
| | PI 442698 | Peru | Cusco | | | C | - |
| | PI 498219 | Peru | Junín | -11.70000000 | -75.43333333 | C | - |
| | PI 568949 | Peru | Puno | | | C | - |
| | PI 310937 | Peru | Cusco | -13.51666667 | -71.98333333 | C | - |
| | PI 458379 | Peru | Apurímac | -13.65000000 | -73.38333333 | C | - |
| | PI 473447 | Peru | Arequipa | -15.86666667 | -74.26666667 | C | - |
| | PI 414155 | Peru | Apurímac | -13.65000000 | -73.38333333 | C | - |
| | PI 568939 | Peru | Apurímac | | | C | - |
| | PI 210042 | Peru | Junín | -11.46666667 | -75.93333333 | C | - |
| | PI 473452 | Peru | | | | C | - |
| | PI 473453 | Peru | Ayacucho | -14.28333333 | -72.10000000 | C | - |
| | PI 473493 | Peru | Huancavelica | -12.83333333 | -74.53333333 | C | - |
| | PI 265876 | Peru | Ayacucho | | | C | - |
| | PI 275271 | Peru | Huánuco | -9.91666667 | -76.23333333 | C | - |
| | PI 210051 | Peru | Junín | -12.68333333 | -75.50000000 | C | - |
| | PI 568933 | Peru | Puno | | | C | - |
| | PI 283084 | Peru | Puno | -15.83333333 | -70.03333333 | S | - |
| | PI 365318 | Peru | Huánuco | -9.91666667 | -76.23333333 | S | - |
| | PI 473491 | Peru | Cusco | -13.30000000 | -71.66666667 | S | - |
| | PI 265865 | Bolivia | La Paz | -17.36694444 | -67.15000000 | W | + |
| *Solanum canasense Hawkes (can)* | PI 246533 | Peru | Cusco | -13.46666667 | -71.91666667 | C | - |
| | PI 265864 | Peru | Cusco | -13.51666667 | -71.98333333 | C | - |
| | PI 265875 | Peru | Cusco | -13.43333333 | -71.85000000 | C | - |
| | PI 458377 | Peru | Puno | -15.83333333 | -70.03333333 | C | - |
| | PI 473346 | Peru | Puno | -15.83333333 | -70.03333333 | C | - |
| | PI 473347 | Peru | Cusco | -14.33333333 | -71.55000000 | C | - |
| | PI 310938 | Peru | Cusco | -13.51666667 | -71.98333333 | C | - |
| | PI 283080 | Peru | Cusco | -13.31666667 | -72.03333333 | C | - |
| | PI 473348 | Peru | Cusco | -13.38333333 | -71.90000000 | C | - |
| | PI 310956 | Peru | Puno | | | S | - |
| | PI 442695 | Peru | Puno | -15.83333333 | -70.03333333 | S | - |
| | PI 458375 | Peru | Puno | -15.83333333 | -70.03333333 | S | - |
| | PI 458376 | Peru | Puno | -15.83333333 | -70.03333333 | S | - |
| | PI 473345 | Peru | Puno | -15.83333333 | -70.03333333 | S | - |
| | PI 265863 | Peru | Puno | -15.83333333 | -70.03333333 | S | + |
| *Solanum coelestipetalum Vargas (cop)* | PI 590904 | Peru | Cusco | | | C | - |
| *Solanum dolichocremastrum Bitt. (dcm)* | PI 498234 | Peru | Ancash | -7.46666667 | -76.61666667 | C | - |
| *Solanum immite Dun. (imt)* | PI 498245 | Peru | La Libertad | -8.11666667 | -79.03333333 | W | - |
| | PI 365330 | Peru | Ancash | -10.16666667 | -77.66666667 | W | - |
| *Solanum marinasense Vargas (mrn)* | PI 310946 | Peru | Puno | -13.63333333 | -71.66666667 | C | + |
| *Solanum medians Bitt. (med)* | PI 210045 | Peru | Lima | -11.41666667 | -76.63333333 | C | - |
| | PI 442703 | Peru | | | | C | - |
| | PI 473496 | Peru | Huánuco | -11.56666667 | -76.61666667 | C | - |
| *Solanum multidissectum Hawkes (mlt)* | PI 210052 | Peru | Ayacucho | -13.26666667 | -73.85000000 | C | - |
| | PI 473349 | Peru | Cusco | -13.43333333 | -71.85000000 | C | - |

| | | | | | | | |
|---|---|---|---|---|---|---|---|
| | PI 210043 | Peru | Junín | -11.45000000 | -75.96666667 | C | - |
| | PI 210044 | Peru | Junín | -11.75000000 | -75.46666667 | C | - |
| | PI 310955 | Peru | Puno | | | S | - |
| | PI 498304 | Peru | Cusco | -14.46666667 | -71.06666667 | S | - |
| | PI 210055 | Peru | Cusco | -14.16666667 | -71.10000000 | S | - |
| | PI 473353 | Peru | Cusco | -14.33333333 | -71.55000000 | S | - |
| *Solanum multiinterruptum Bitt. (mtp)* | PI 275272 | Peru | Cusco | -13.33333333 | -71.96666667 | C | - |
| *Solanum pampasense Hawkes (pam)* | PI 275274 | Peru | Ayacucho | -13.45000000 | -73.73333333 | W | - |
| *Solanum chaucha Juz. et Buk. (cha)* | CIP 701013 | Peru | Cajamarca | -6.92390000 | -78.13430000 | A | - |
| *Solanum chaucha Juz. et Buk. (cha)* | CIP 702551 | | | | | S | - |
| *Solanum juzepczukii Buk. (juz)* | CIP 700895 | | | | | C | - |
| *Solanum phureja Juz. et Buk. (phu)* | CIP 703272 | Bolivia | La Paz | -17.45000000 | -67.51660000 | A | - |
| | CIP 703291 | Colombia | Cauca | 2.51380000 | -76.39580000 | S | - |
| | CIP 703293 | Colombia | Cauca | 2.35000000 | -76.50000000 | S | - |
| | CIP 703274 | Peru | Ayacucho | -12.93770000 | -74.24870000 | S | - |
| | CIP 703308 | Peru | Lambayeque | -6.23460000 | -79.31700000 | S | - |
| | CIP 703275 | Peru | Puno | -14.15230000 | -69.66510000 | S | - |
| *Solanum stenotomum Juz. et Buk. (stn)* | CIP 707297 | Peru | Cusco | -13.38000000 | -71.67000000 | S | - |
| | CIP 704141 | Bolivia | 0 | | | A | - |
| | CIP 703761 | Bolivia | La Paz | -17.33330000 | -67.73330000 | A | - |
| | CIP 701960 | Peru | Pasco | -10.63680000 | -75.95010000 | A | - |
| | CIP 703871 | Peru | Huánuco | -9.87420000 | -76.81200000 | S | - |
| | CIP 703823 | Peru | Pasco | -10.43860000 | -76.49470000 | S | - |
| | CIP 704065 | Bolivia | Oruro | -17.76660000 | -67.48330000 | S | - |
| | CIP 703475 | Bolivia | Chuquisaca | -19.04300000 | -65.25910000 | S | - |
| | CIP 703808 | Peru | Puno | -16.27290000 | -69.29310000 | S | - |
| | CIP 703710 | Peru | Junín | -10.96670000 | -75.87740000 | S | - |
| | CIP 703933 | Peru | Cusco | -13.24760000 | -71.89690000 | A | - |
| | CIP 705980 | Bolivia | La Paz | -17.01660000 | -68.61660000 | S | - |
| | CIP 703625 | Peru | Cusco | -13.45720000 | -72.25720000 | S | - |
| | CIP 703983 | Peru | Apurímac | -13.56920000 | -73.34430000 | S | - |
| | CIP 704969 | Bolivia | La Paz | | | S | - |
| | CIP 703712 | Peru | Junín | -10.96670000 | -75.87740000 | S | - |
| | CIP 704330 | | | | | S | - |
| | CIP 704014 | Peru | Cusco | -16.48000000 | -69.10000000 | S | - |
| | CIP 703959 | Peru | Cusco | -14.62660000 | -71.53140000 | S | - |
| | CIP 700348 | Peru | Junín | -12.13850000 | -75.25680000 | S | - |
| | CIP 705638 | Peru | Junín | -11.75040000 | -75.13200000 | S | - |
| | CIP 704438 | Peru | Pasco | -10.63680000 | -75.95010000 | S | - |
| | CIP 701243 | Peru | Ancash | -10.05150000 | -77.13590000 | A | - |
| | CIP 706846 | | | | | A | - |
| | CIP 706233 | Bolivia | Cochabamba | -17.43330000 | -65.71660000 | S | - |
| | CIP 704917 | Bolivia | Cochabamba | -17.43330000 | -65.71660000 | S | - |
| | CIP 704950 | Bolivia | Cochabamba | -17.43330000 | -65.71660000 | S | - |
| | CIP 706190 | Peru | Ayacucho | -13.23000000 | -74.35000000 | A | - |
| | CIP 705476 | Peru | Ayacucho | -12.95000000 | -74.02000000 | S | - |
| | CIP 703624 | Peru | Puno | -11.08790000 | -76.14750000 | S | - |
| | CIP 703996 | Peru | Cusco | -13.21910000 | -73.02200000 | S | - |
| | CIP 704090 | Bolivia | Potosi | -10.43860000 | -76.49470000 | S | - |
| | CIP 704827 | Bolivia | Oruro | -18.32120000 | -67.68330000 | S | - |
| | CIP 703421 | Bolivia | La Paz | -16.90000000 | -68.10000000 | A | - |
| | CIP 704801 | Bolivia | Oruro | -17.98330000 | -67.85000000 | A | - |
| | CIP 705961 | Bolivia | La Paz | -17.50000000 | -68.50000000 | S | - |
| | CIP 704796 | Bolivia | Oruro | -18.35000000 | -67.93330000 | S | - |
| | CIP 705962 | Bolivia | La Paz | -17.40000000 | -69.06660000 | A | - |
| | CIP 704771 | Bolivia | Oruro | -16.70000000 | -68.08330000 | A | - |
| | CIP 705534 | Peru | Apurímac | -13.69550000 | -73.15210000 | S | - |
| | CIP 705861 | Peru | Ayacucho | -13.23000000 | -74.35000000 | S | - |
| | CIP 704956 | Bolivia | Cochabamba | -17.43330000 | -65.71660000 | S | - |
| | CIP 704862 | Bolivia | Cochabamba | -17.43330000 | -65.71660000 | S | - |
| | CIP 706250 | Bolivia | Cochabamba | -17.43330000 | -65.71660000 | A | - |
| | CIP 704890 | Bolivia | Cochabamba | -17.43330000 | -65.71660000 | S | - |
| | CIP 706845 | Bolivia | Cochabamba | -17.43330000 | -65.71660000 | S | - |
| | CIP 704805 | Bolivia | Oruro | -17.90000000 | -67.80000000 | S | - |
| | CIP 704819 | Bolivia | La Paz | -17.50000000 | -68.50000000 | A | - |
| | CIP 704970 | Bolivia | Potosi | -19.83000000 | -65.50000000 | A | - |
| | CIP 702588 | Bolivia | La Paz | -17.81660000 | -69.06660000 | S | - |
| | CIP 704571 | Peru | Apurímac | -14.31310000 | -72.94420000 | S | - |
| | CIP 704830 | Bolivia | Oruro | -18.16660000 | -67.41660000 | S | - |
| | CIP 704834 | Bolivia | Cochabamba | -17.43330000 | -65.71660000 | S | - |
| | CIP 704914 | Bolivia | Cochabamba | -17.43330000 | -65.71660000 | S | - |
| | CIP 705940 | Peru | Junín | -12.27820000 | -75.14840000 | A | - |
| | CIP 705839 | Peru | Cusco | -13.36000000 | -71.67340000 | A | - |
| | CIP 702172 | Peru | Junín | -12.13850000 | -75.25680000 | A | - |
| | CIP 701676 | Peru | Junín | -12.12370000 | -75.43750000 | S | - |
| | CIP 702815 | Peru | Puno | -15.83920000 | -70.02920000 | S | - |
| | CIP 703088 | Peru | Junín | -11.26490000 | -75.65030000 | C | - |
| | CIP 703276 | Peru | Puno | -16.90870000 | -69.37080000 | S | - |
| | CIP 703287 | Peru | Cusco | -13.47970000 | -71.92230000 | S | - |
| | CIP 703312 | Peru | Puno | -14.47120000 | -69.53740000 | A | - |

| | | | | | | | |
|---|---|---|---|---|---|---|---|
| | CIP 703314 | Peru | Puno | -14.45470000 | -69.53710000 | S | - |
| | CIP 703396 | Peru | Ancash | -9.18740000 | -76.99260000 | S | - |
| | CIP 702580 | Bolivia | Oruro | -17.53330000 | -67.23330000 | S | - |
| | CIP 703470 | Peru | Ayacucho | -13.07700000 | -73.74520000 | S | - |
| | CIP 703319 | Peru | Puno | -15.83920000 | -70.02920000 | S | - |
| | CIP 700407 | Peru | Cusco | -13.45720000 | -72.25720000 | S | - |
| | CIP 701633 | Peru | Junín | -12.13850000 | -75.25680000 | S | - |
| | CIP 702286 | Bolivia | Cochabamba | -17.43330000 | -65.71660000 | S | - |
| | CIP 702421 | Peru | Cusco | -13.31850000 | -71.59550000 | S | - |
| | CIP 702583 | Bolivia | Potosi | -19.63330000 | -65.86660000 | W | - |
| | CIP 702834 | Peru | Puno | -15.79930000 | -70.02140000 | S | - |
| | CIP 703698 | Peru | Junín | -11.75040000 | -75.13200000 | S | - |
| | CIP 703709 | Peru | Junín | -11.08790000 | -76.14750000 | A | - |
| | CIP 703317 | Peru | Puno | -9.57350000 | -77.51530000 | S | - |
| | CIP 703473 | Bolivia | La Paz | -16.50000000 | -68.15000000 | A | - |
| | CIP 702547 | Bolivia | Potosi | -19.21830000 | -65.82630000 | S | - |
| | CIP 703313 | Peru | Junín | -11.70000000 | -75.03000000 | A | - |
| | CIP 701165 | Peru | Junín | -12.13850000 | -75.25680000 | S | - |
| | CIP 701985 | Peru | Cusco | -13.45720000 | -72.25720000 | S | - |
| | CIP 702199 | Peru | Cusco | -13.43270000 | -71.50280000 | S | - |
| | CIP 700235 | Peru | Puno | -16.24590000 | -69.09180000 | S | - |
| | CIP 703286 | Bolivia | La Paz | -17.33330000 | -67.73330000 | S | - |
| *Solanum tuberosum L.* ssp. *andigena Hawkes (adg)* | PI 161350 | Mexico | Puebla | 19.01666667 | -97.26666667 | A | - |
| | PI 281080 | Peru | Cusco | -13.51666667 | -71.98333333 | A | - |
| | PI 281093 | Peru | Cusco | -13.51666667 | -71.98333333 | A | - |
| | PI 292078 | Peru | Arequipa | -16.40000000 | -71.55000000 | C | - |
| | PI 292089 | Peru | La Libertad | -8.11666667 | -79.03333333 | C | - |
| | PI 214426 | Peru | Huánuco | -9.91666667 | -76.23333333 | A | - |
| | PI 292096 | Peru | Cajamarca | -7.16666667 | -78.51666667 | C | - |
| | PI 292097 | Peru | Cajamarca | -7.16666667 | -78.51666667 | C | - |
| | PI 292107 | Peru | Huánuco | -9.91666667 | -76.23333333 | A | - |
| | PI 365345 | Peru | La Libertad | -8.28333333 | -77.30000000 | W | - |
| | PI 365372 | Peru | Ayacucho | -12.93333333 | -74.25000000 | W | + |
| | PI 214429 | Peru | Pasco | -10.68333333 | -76.26666667 | A | - |
| | PI 473196 | Argentina | Jujuy | -22.11666667 | -65.46666667 | A | - |
| | PI 473246 | Argentina | Salta | -24.68333333 | -65.75000000 | A | - |
| | PI 473249 | Argentina | Jujuy | -22.13333333 | -65.75000000 | A | - |
| | PI 473251 | Argentina | Jujuy | -21.88333333 | -66.18333333 | A | - |
| | PI 473253 | Argentina | Jujuy | -21.88333333 | -66.18333333 | A | - |
| | PI 473254 | Argentina | Jujuy | -22.43333333 | -66.11666667 | W | + |
| | PI 473255 | Argentina | Jujuy | -22.43333333 | -66.11666667 | A | - |
| | PI 473257 | Argentina | Salta | -24.65000000 | -66.20000000 | T | + |
| | PI 214430 | Peru | Junín | -11.53333333 | -75.90000000 | A | - |
| | PI 473260 | Argentina | Salta | -22.15000000 | -65.03333333 | S | - |
| | PI 473261 | Argentina | Salta | -22.15000000 | -65.03333333 | S | - |
| | PI 473262 | Argentina | Salta | -22.15000000 | -65.03333333 | S | - |
| | PI 473265 | Argentina | Jujuy | -23.58333333 | -65.33333333 | A | - |
| | PI 473267 | Argentina | Jujuy | -23.16666667 | -65.45000000 | A | - |
| | PI 473268 | Argentina | Salta | -24.36666667 | -66.06666667 | W | + |
| | PI 473269 | Argentina | Salta | -24.36666667 | -66.06666667 | A | - |
| | PI 473270 | Argentina | Salta | -23.93333333 | -66.35000000 | A | - |
| | PI 214434 | Peru | Junín | -11.53333333 | -75.90000000 | A | - |
| | PI 473275 | Argentina | Jujuy | -22.11666667 | -65.46666667 | A | - |
| | PI 473276 | Argentina | Jujuy | -22.13333333 | -65.71666667 | A | - |
| | PI 473277 | Argentina | Jujuy | -22.11666667 | -65.70000000 | A | - |
| | PI 473278 | Argentina | Jujuy | -21.88333333 | -66.18333333 | C | - |
| | PI 473281 | Argentina | Jujuy | -23.86666667 | -65.80000000 | W | + |
| | PI 473282 | Argentina | Salta | -23.93333333 | -66.35000000 | A | - |
| | PI 473283 | Argentina | Salta | -23.93333333 | -66.35000000 | A | - |
| | PI 473284 | Argentina | Salta | -24.31666667 | -66.10000000 | A | - |
| | PI 473285 | Argentina | Salta | -24.93333333 | -66.16666667 | W | + |
| | PI 473286 | Argentina | Salta | -24.93333333 | -66.16666667 | A | - |
| | PI 214436 | Peru | Junín | -11.53333333 | -75.90000000 | A | - |
| | PI 473287 | Argentina | Salta | -24.71666667 | -66.20000000 | A | - |
| | PI 473288 | Argentina | Salta | -24.71666667 | -66.20000000 | A | - |
| | PI 473290 | Argentina | Salta | -25.21666667 | -66.21666667 | A | - |
| | PI 473291 | Argentina | Salta | -25.05000000 | -66.21666667 | A | - |
| | PI 473292 | Argentina | Jujuy | -22.10000000 | -65.95000000 | A | - |
| | PI 473293 | Argentina | Jujuy | -22.10000000 | -65.46666667 | C | - |
| | PI 473294 | Argentina | Jujuy | -22.11666667 | -65.46666667 | C | - |
| | PI 473295 | Argentina | Jujuy | -22.11666667 | -65.46666667 | C | - |
| | PI 473296 | Argentina | Jujuy | -22.11666667 | -65.46666667 | A | - |
| | PI 473298 | Argentina | Jujuy | -22.11666667 | -65.46666667 | C | - |
| | PI 214441 | Peru | Lima | -12.05000000 | -77.05000000 | A | - |
| | PI 473299 | Argentina | Salta | -22.26666667 | -64.93333333 | C | - |
| | PI 473300 | Argentina | Salta | -22.36666667 | -65.06666667 | A | - |
| | PI 473301 | Argentina | Salta | -22.40000000 | -65.11666667 | A | - |
| | PI 473302 | Argentina | Salta | -22.40000000 | -65.11666667 | C | - |
| | PI 473390 | Bolivia | Potosí | -19.63333333 | -65.06666667 | C | - |
| | PI 473391 | Bolivia | Cochabamba | -17.68333333 | -66.60000000 | W | + |

| | | | | | | |
|---|---|---|---|---|---|---|
| PI 473507 | Bolivia | Chuquisaca | -21.33333333 | -66.23333333 | C | - |
| PI 473508 | Bolivia | Chuquisaca | -19.15000000 | -64.91666667 | A | - |
| PI 498076 | Bolivia | Chuquisaca | -19.10000000 | -65.23333333 | W | - |
| PI 214442 | Peru | Lima | -12.05000000 | -77.05000000 | A | - |
| PI 498291 | Bolivia | Chuquisaca | -19.00000000 | -65.26666667 | A | - |
| PI 498294 | Bolivia | Chuquisaca | -20.91666667 | -64.91666667 | C | - |
| PI 498307 | Bolivia | Chuquisaca | -20.25000000 | -65.18333333 | C | - |
| PI 498309 | Bolivia | Potosí | -20.93333333 | -65.63333333 | A | - |
| PI 498310 | Bolivia | La Paz | 0.00000000 | | S | - |
| PI 500057 | Argentina | Jujuy | -24.00000000 | -65.61666667 | C | - |
| PI 500058 | Argentina | Jujuy | -24.08333333 | -65.65000000 | A | - |
| PI 214443 | Peru | Lima | -12.05000000 | -77.05000000 | A | - |
| PI 545930 | Bolivia | Cochabamba | -17.43333333 | -65.46666667 | C | - |
| PI 546016 | Bolivia | Cochabamba | -17.61666667 | -66.71666667 | A | + |
| PI 546017 | Bolivia | La Paz | -15.63333333 | -69.06666667 | W | - |
| PI 546018 | Bolivia | La Paz | -15.73333333 | -68.96666667 | S | - |
| PI 546020 | Bolivia | Cochabamba | -17.93333333 | -66.46666667 | C | - |
| PI 546021 | Bolivia | Potosí | -18.01666667 | -66.36666667 | S | - |
| PI 546023 | Bolivia | Potosí | -18.01666667 | -66.38333333 | A | - |
| PI 546024 | Bolivia | Cochabamba | -17.63333333 | -66.71666667 | C | - |
| PI 546025 | Bolivia | Cochabamba | -18.03333000 | -64.86667000 | W | - |
| PI 546026 | Bolivia | Chuquisaca | -18.93333333 | -65.38333333 | C | - |
| PI 225628 | Colombia | Santander | 7.13333333 | -73.15000000 | A | - |
| PI 546028 | Bolivia | Chuquisaca | -19.45000000 | -64.88333333 | W | - |
| PI 558137 | Argentina | Salta | -25.06666667 | -66.03333333 | A | - |
| PI 558139 | Argentina | Jujuy | -23.50000000 | -65.45000000 | S | - |
| PI 558141 | Argentina | Jujuy | -23.60000000 | -65.58333333 | T | + |
| PI 558142 | Argentina | Jujuy | -24.10000000 | -65.66666667 | A | - |
| PI 558144 | Argentina | Jujuy | -23.16666667 | -65.18333333 | A | - |
| PI 558145 | Argentina | Salta | -24.65000000 | -66.20000000 | A | - |
| PI 473393 | Bolivia | Potosí | -19.56666667 | -65.43333333 | W | - |
| PI 161683 | Mexico | Tlaxcala | 19.61666667 | -98.11666667 | A | - |
| PI 225633 | Colombia | Boyacá | 5.48333333 | -73.26666667 | A | - |
| PI 225635 | Venezuela | Mérida | 8.75000000 | -70.91666667 | A | - |
| PI 230496 | Peru | Tacna | -17.50000000 | -70.03333333 | S | - |
| PI 230497 | Peru | Tacna | -17.50000000 | -70.03333333 | A | - |
| PI 230499 | Peru | Tacna | -17.50000000 | -70.03333333 | C | - |
| PI 232036 | Peru | Pasco | -11.85000000 | -77.03333333 | C | - |
| PI 232839 | Peru | Pasco | -11.85000000 | -77.03333333 | A | - |
| PI 232840 | Peru | Cusco | -13.51666667 | -71.98333333 | C | - |
| PI 232841 | Peru | Pasco | -12.75000000 | -76.33333333 | S | - |
| PI 161716 | Mexico | Michoacán de Ocampo | 20.76666667 | -101.33333333 | A | - |
| PI 237208 | Ecuador | Cotopaxi | -0.93333333 | -78.61666667 | A | - |
| PI 243360 | Colombia | Cundinamarca | 5.03333333 | -74.00000000 | A | - |
| PI 243363 | Colombia | Boyacá | 5.78333333 | -73.25000000 | A | - |
| PI 243400 | Ecuador | Carchi | 0.60000000 | -77.81666667 | A | - |
| PI 243401 | Ecuador | Carchi | 0.60000000 | -77.81666667 | A | - |
| PI 243405 | Ecuador | Imbabura | 0.30000000 | -78.58333333 | A | - |
| PI 184903 | Guatemala | San Marcos | 15.03277778 | -91.79805556 | A | - |
| PI 243406 | Ecuador | Imbabura | 0.30000000 | -78.58333333 | A | - |
| PI 243407 | Ecuador | Imbabura | 0.30000000 | -78.58333333 | A | - |
| PI 243409 | Ecuador | Imbabura | 0.30000000 | -78.58333333 | A | - |
| PI 243411 | Ecuador | Imbabura | 0.30000000 | -78.58333333 | A | - |
| PI 243415 | Colombia | Cauca | 2.63333333 | -76.50000000 | A | - |
| PI 243429 | Colombia | Caldas | 4.63333333 | -75.53333333 | A | - |
| PI 243430 | Colombia | Tolima | 4.43333333 | -75.45000000 | A | - |
| PI 243431 | Colombia | Antioquia | 6.15000000 | -75.36666667 | A | - |
| PI 243434 | Colombia | Santander | 7.13333333 | -73.15000000 | A | - |
| PI 186178 | Peru | Huánuco | -12.01666667 | -75.25000000 | C | - |
| PI 243436 | Colombia | Norte de Santander | 7.38333333 | -72.65000000 | A | - |
| PI 243441 | Colombia | Nariño | 0.83333333 | -77.61666667 | A | - |
| PI 246497 | Peru | Ancash | -9.56666667 | -78.20000000 | C | - |
| PI 246499 | Peru | Ancash | -9.56666667 | -78.20000000 | A | - |
| PI 246516 | Peru | Puno | -15.83333333 | -70.03333333 | A | - |
| PI 246521 | Peru | Puno | -15.83333333 | -70.03333333 | A | - |
| PI 246545 | Peru | Cusco | -13.43333333 | -71.85000000 | A | - |
| PI 186179 | Peru | Huánuco | -12.01666667 | -75.25000000 | A | - |
| PI 246555 | Peru | Cusco | -13.43333333 | -71.85000000 | A | - |
| PI 255503 | Argentina | Salta | -25.16666667 | -65.76666667 | A | - |
| PI 255505 | Bolivia | La Paz | -15.80000000 | -69.40000000 | A | - |
| PI 255508 | Bolivia | Chuquisaca | -21.45000000 | -65.71666667 | W | - |
| PI 258857 | Bolivia | La Paz | -15.50000000 | -68.00000000 | C | - |
| PI 258879 | Bolivia | Oruro | -18.90000000 | -66.78333333 | C | - |
| PI 258881 | Bolivia | Oruro | -15.88333333 | -68.11666667 | C | - |
| PI 258886 | Bolivia | Cochabamba | -18.35000000 | -68.95000000 | W | + |
| PI 195162 | Peru | Huánuco | -9.91666667 | -76.23333333 | C | - |
| PI 258917 | Bolivia | Cochabamba | -16.93333333 | -66.70000000 | C | - |
| PI 258927 | Bolivia | Cochabamba | -17.35000000 | -65.86666667 | A | - |
| PI 258936 | Bolivia | La Paz | -15.58333333 | -68.71666667 | C | - |
| PI 265882 | Bolivia | Chuquisaca | -19.03333333 | -65.28333333 | W | - |

| | | | | | | | |
|---|---|---|---|---|---|---|---|
| | PI 275119 | Bolivia | Oruro | -15.38333333 | -67.90000000 | S | - |
| | PI 280989 | Bolivia | La Paz | -16.55000000 | -68.70000000 | A | - |
| | PI 280990 | Bolivia | La Paz | -16.58333333 | -68.81666667 | A | - |
| | PI 197757 | Bolivia | Tarija | | | A | - |
| | PI 280993 | Bolivia | La Paz | -17.48333333 | -66.16666667 | A | - |
| | PI 281008 | Bolivia | La Paz | -15.53333333 | -69.25000000 | A | - |
| | PI 281021 | Bolivia | Oruro | -15.38333333 | -67.90000000 | C | - |
| | PI 281031 | Bolivia | Oruro | -15.38333333 | -67.90000000 | C | - |
| | PI 281032 | Bolivia | Oruro | -15.38333333 | -67.90000000 | A | - |
| | PI 281059 | Peru | Huánuco | -11.41666667 | -75.70000000 | A | - |
| | PI 281060 | Peru | Huánuco | -11.41666667 | -75.70000000 | A | - |
| | PI 281061 | Peru | Cusco | -13.40000000 | -72.05000000 | C | - |
| | PI 281064 | Peru | Cusco | -13.40000000 | -72.05000000 | A | - |
| | PI 281066 | Peru | Cusco | -13.40000000 | -72.05000000 | A | - |
| | CIP 704111 | Venezuela | Trujillo | 9.36660000 | -70.43330000 | A | - |
| | CIP 700921 | Peru | Cusco | -13.60480000 | -71.56110000 | C | - |
| | CIP 701624 | Peru | Junín | -12.13850000 | -75.25680000 | C | - |
| | CIP 702453 | Peru | Puno | -16.08590000 | -69.63810000 | A | - |
| | CIP 700017 | Peru | Junín | -12.13850000 | -75.25680000 | A | - |
| | CIP 700045 | Peru | Junín | -12.13850000 | -75.25680000 | A | - |
| | CIP 700094 | Peru | Junín | -12.13850000 | -75.25680000 | A | - |
| | CIP 700387 | Peru | Cusco | -13.51410000 | -72.06980000 | S | - |
| | CIP 703268 | Ecuador | Quito | -0.36660000 | -78.51660000 | A | - |
| | CIP 700532 | Peru | Ancash | -8.53450000 | -77.56630000 | A | - |
| | CIP 700598 | Peru | Junín | -10.96670000 | -75.87740000 | C | - |
| | CIP 700616 | Peru | Junín | -12.13850000 | -75.25680000 | A | - |
| | CIP 700652 | Peru | Junín | -12.13850000 | -75.25680000 | C | - |
| | CIP 700696 | Peru | Junín | -12.13850000 | -75.25680000 | A | - |
| | CIP 700767 | Peru | Cusco | -13.45720000 | -72.25720000 | A | - |
| | CIP 700771 | Peru | Ancash | -8.53030000 | -77.48750000 | S | - |
| | CIP 700787 | Peru | Junín | -12.13850000 | -75.25680000 | C | - |
| | CIP 700790 | Peru | Puno | -15.29670000 | -69.97810000 | A | - |
| | CIP 701304 | | | | | A | - |
| | CIP 700863 | Peru | Cusco | -13.60480000 | -71.56110000 | C | - |
| | CIP 700877 | Peru | La Libertad | -11.82000000 | -75.38000000 | C | - |
| | CIP 700882 | Peru | Junín | -12.13850000 | -75.25680000 | A | - |
| | CIP 700960 | Peru | Junín | -12.13850000 | -75.25680000 | S | - |
| | CIP 701201 | Peru | Junín | -12.13850000 | -75.25680000 | S | - |
| | CIP 702535 | Bolivia | Potosi | -19.21830000 | -65.82630000 | A | - |
| | CIP 702698 | Bolivia | Potosi | -20.95000000 | -68.03330000 | A | - |
| | CIP 703474 | Bolivia | Potosi | -19.63330000 | -65.06660000 | A | - |
| | CIP 704082 | Bolivia | Potosi | -18.93330000 | -65.80000000 | S | - |
| | CIP 701306 | Peru | Junín | -12.13850000 | -75.25680000 | S | - |
| | CIP 701065 | Peru | Junín | -12.13850000 | -75.25680000 | A | - |
| | CIP 701067 | Peru | Junín | -12.13850000 | -75.25680000 | A | - |
| | CIP 701074 | | | | | S | - |
| | CIP 704152 | Argentina | Salta | -22.43330000 | -64.81660000 | A | - |
| | CIP 703653 | Peru | Cajamarca | -6.91090000 | -78.25480000 | A | - |
| | CIP 701296 | Peru | Ancash | -8.51240000 | -77.86480000 | A | - |
| | CIP 701463 | Peru | Junín | -11.47130000 | -75.68750000 | A | - |
| | CIP 703682 | Peru | Huancavelica | -12.79010000 | -74.58550000 | A | - |
| *Solanum tuberosum L.* ssp. *tuberosum (tbr)* | CIP 704165 | Chile | Archp. Los Chonos | -42.66660000 | -73.91660000 | A | - |
| | CIP 703252 | Chile | Archp. Los Chonos | -42.66660000 | -73.91660000 | T | + |
| | CIP 703254 | Chile | Archp. Los Chonos | -43.86330000 | -73.99330000 | T | + |
| | CIP 703610 | Chile | Archp. Los Chonos | -43.86330000 | -73.99330000 | T | + |
| | CIP 704168 | Chile | Archp. Los Chonos | -42.66660000 | -73.91660000 | T | + |
| | CIP 704171 | Chile | Archp. Los Chonos | -42.66660000 | -73.91660000 | T | + |
| | CIP 704172 | Chile | Archp. Los Chonos | -42.66660000 | -73.91660000 | T | + |

+ indicates that the gene was confirmed to be present by PCR.
- indicates that the gene was confirmed to be absent by PCR.
Blank cells indicate that data were not available for that entry.
* The classification system of potato follows Hawkes (1990).

**Supplemental Table 3** Primers used in this study

| Name | Sequence (5'-3') | Application |
|---|---|---|
| tomato_orf320_Fw | AAAAATAAATTCCTTTTTTTGAGCTCCCGA | RT-PCR for tomato *orf320* |
| tomato_orf320_Rv | TCAAAACTTAAGAATCTCTTCGACTATCAT | RT-PCR for tomato orf320 |
| potato_orf320_Fw | CGATTGCTCGACAGGACTTAGA | PCR and RT-PCR for potato *orf320* |
| potato_orf320_Rv | CCTTCCGCGGAATATATGACTCA | PCR and RT-PCR for potato orf320 |
| cox2_Fw | CCCGCAAAGGATTGTTCATGG | RT-PCR for tomato and potato *cox2* |
| cox2_Rv | CGTATAGGGCTCTTTGCTGGTAG | RT-PCR for tomato and potato cox2 |
| orf137_Fw | CGATTGAGAAAGCGGCAGGC | PCR for potato *orf137* |
| orf137_Rv | GTTATTTTCGCTGCAACGGCG | PCR for potato *orf137* |
| NPTII_Fw | ATGATTGAACAAGATGGATTGCAC | PCR for *NPTII* |
| NPTII_Rv | TCAGAAGAACTCGTCAAGAAGGCG | PCR for *NPTII* |
