## Supplemental Figure for "Relationship Between Open Reading Frame 320, a Gene Causing Male Sterility in Tomatoes, and Cytoplasmic Male Sterility in Potatoes"

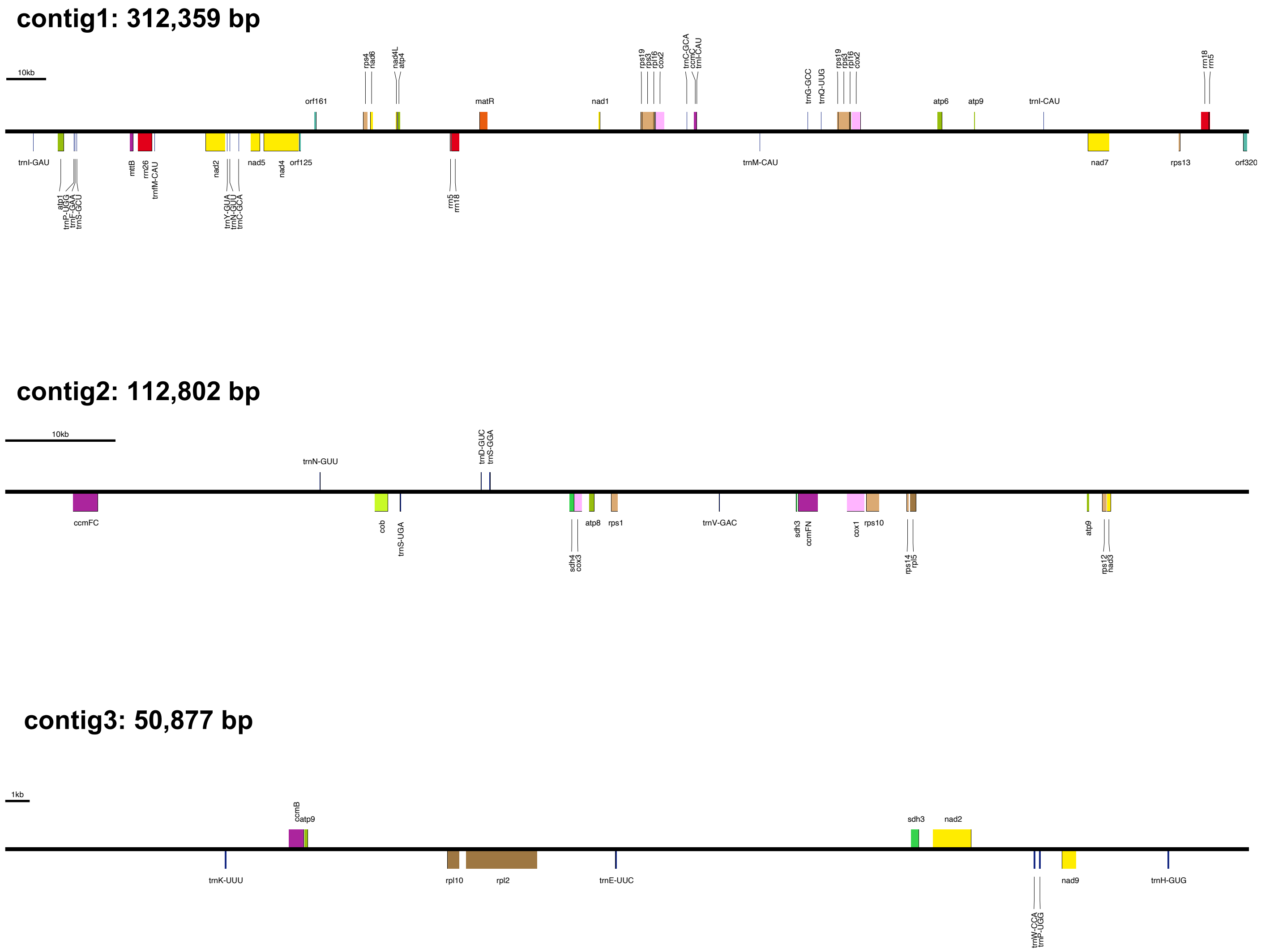


**Supplementary Fig. S1** Gene annotations for the mitochondrial genome of May Queen. The assembled genome was annotated using the online tool PMGA with Dataset 2 (Li et al. 2024) and visualized using OGDRAW (Greiner et al. 2019).

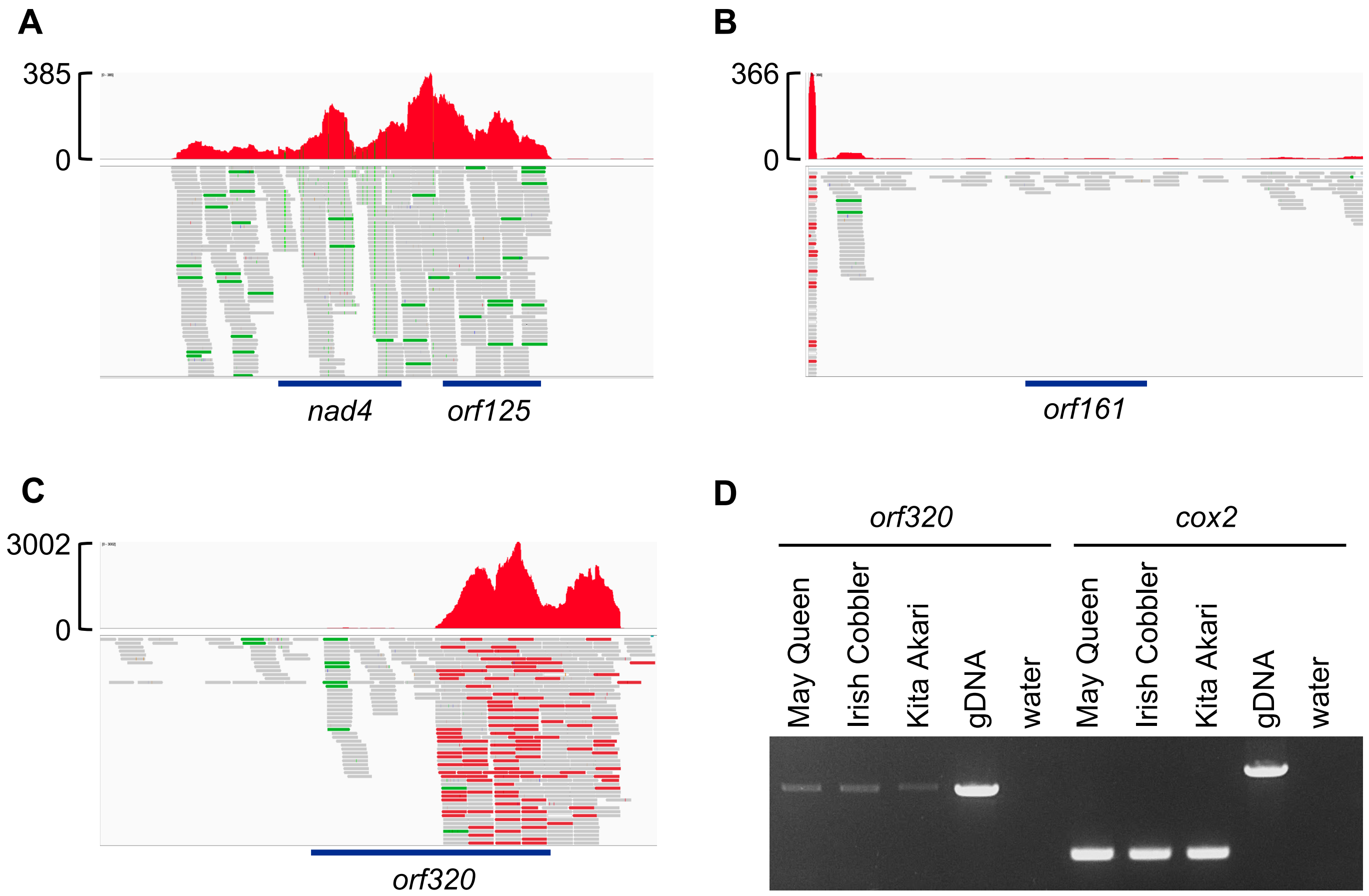


**Supplementary Fig. S2** Expression analysis for *orf125*, *orf161*, and *orf320*.
(A–C) RNA-Seq analysis showing the expression levels of mitochondrial genes, *orf125*, *orf161*, and *orf320* in potato anthers. Reads are displayed as horizontal gray, red, and green bars. The upper red peaks represent coverage depth. (D) Expression analysis for *orf320* in various potato cultivars. Amplification of *orf320* was observed in the anthers of T/β cytoplasm varieties, May Queen, Irish Cobbler, and Kita Akari. The *cox2* primers were designed to span an intron, producing a shorter amplicon in cDNA and a longer product in genomic DNA. No long bands were detected in cDNA, meaning no contamination of gDNA.

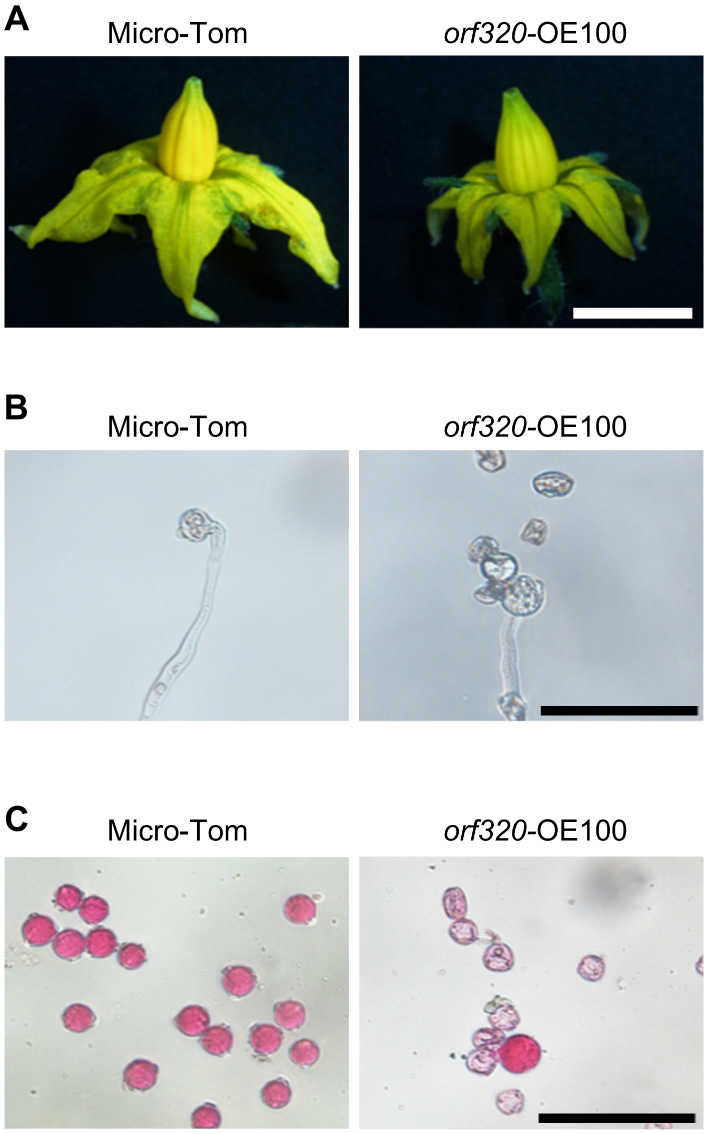


**Supplementary Fig. S3** Phenotypes of flowers and pollen in *orf320*-overexpressed tomatoes. (A) Overexpression of *orf320* without *MTP* showed pale yellow anthers. Scale bar = 5 mm. (B) *orf320*-OE100 exhibited reduced pollen fertility in the *in vitro* pollen germination assay. Scale bar = 100 µm. (C) *orf320*-OE100 exhibited reduced pollen fertility in the Alexander staining assay. Micro-Tom: pollen fertility = 95%, *orf320*-OE100: pollen fertility = 14%. Scale bar = 100 µm.

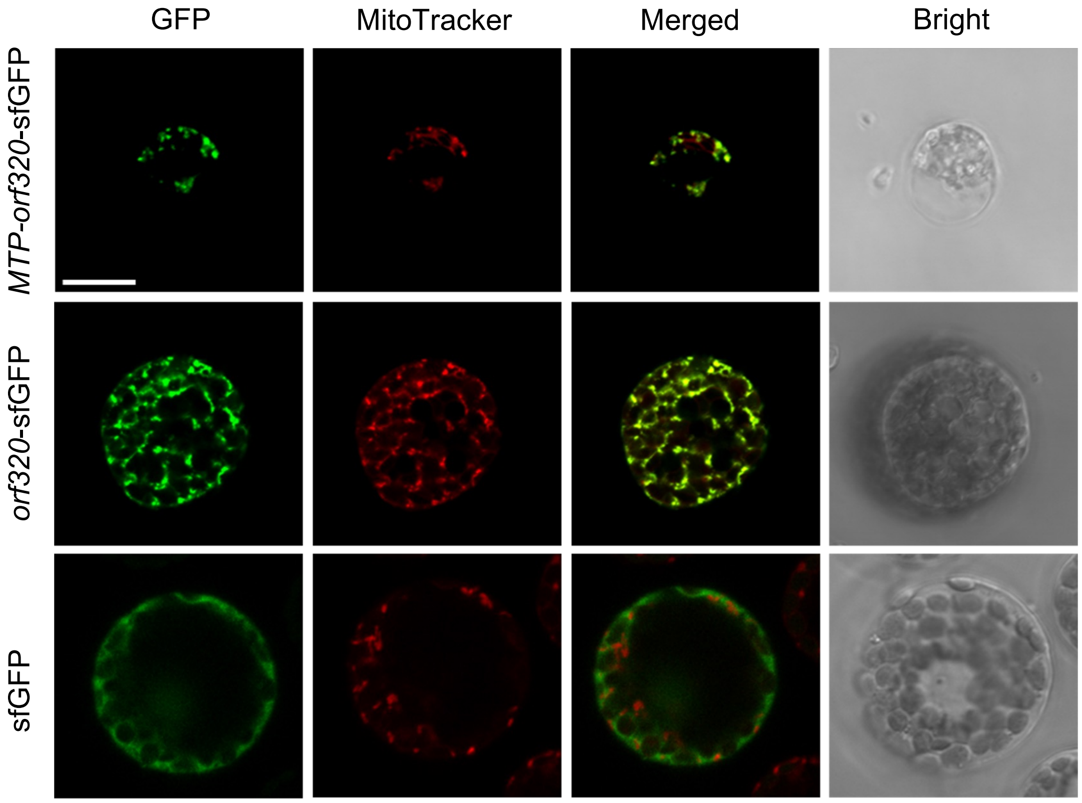


**Supplementary Fig. S4** Subcellular localization of MTP-ORF320 and ORF320 in *Nicotiana benthamiana* protoplasts.
Fluorescent images of protoplasts expressing *MTP-orf320-sfGFP*, *orf320-sfGFP*, and *sfGFP* as a negative control. GFP fluorescence (green) indicates the localization of the fusion proteins, while mitochondria were stained with MitoTracker Red. Merged images show colocalization of GFP signals with mitochondria. Bar = 10 µm.

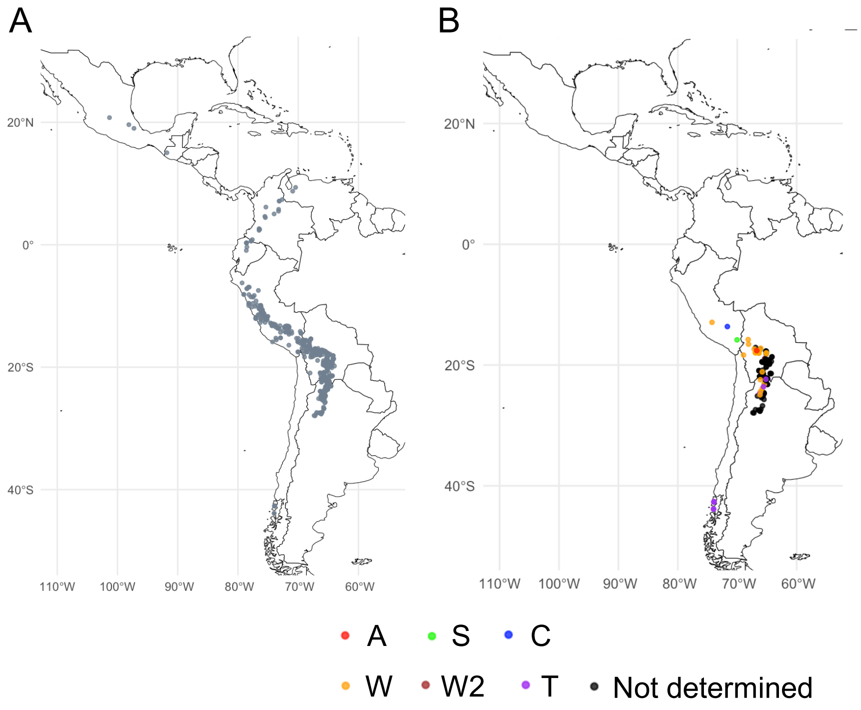


**Supplementary Fig. S5** Geographical distribution of wild potato species and relatives with *orf320*. (A) The geographic distribution of surveyed potato lines. (B) The distribution of potato lines carrying the *orf320* gene. Different color plots indicate the cpDNA types.
